## Supplemental material for "Genomic evidence reveals three sequential W-autosome fusions in *Heliconius* butterflies"

### Tables

**Table S1.** Sample information and genotyping statistics. Individuals in bold were not included in the analyses due to low depth and missing data.

| Sequence ID | taxon | sex | country | latitude | longitude | mean read depth | missing data per indv. | high quality calls, % QUAL >30 | % mapping genome | Accession | Publication |
|---|---|---|---|---|---|---|---|---|---|---|---|
| M4096 | *H. antiochus antiochus* | Female | Colombia | -4.16 | -69.97 | 18.01 | 0.13 | 94.31 | 95.86 | ERR12713877 | This study |
| M4097 | *H. antiochus antiochus* | Male | Colombia | -4.16 | -69.97 | 18.78 | 0.13 | 94.39 | 95.91 | ERR12713878 | This study |
| S21203 | *H. antiochus antiochus* | Female | Brazil | 4.74 | -56.80 | 18.74 | 0.13 | 94.42 | 96.01 | ERR12713879 | This study |
| S21229 | *H. antiochus antiochus* | Male | Brazil | 4.74 | -56.80 | 16.86 | 0.14 | 94.69 | 96.17 | ERR12713880 | This study |
| S21234 | *H. antiochus antiochus* | Female | Brazil | 4.74 | -56.80 | 18.80 | 0.13 | 93.78 | 96.24 | ERR12713881 | This study |
| S337_1 | *H. antiochus antiochus* | Male | Brazil | -13.49 | 61.03 | 18.41 | 0.13 | 92.75 | 96.10 | ERR12713882 | This study |
| S4897 | *H. antiochus aranea* | Male | Colombia | 2.58 | -72.72 | 17.36 | 0.14 | 93.15 | 96.11 | ERR12713883 | This study |
| R5074 | *H. antiochus aranea* | Female | Colombia | 5.34 | -72.42 | 13.20 | 0.15 | 93.35 | 96.94 | ERR12713884 | This study |
| S9556 | *H. antiochus aranea* | Female | Panama | 8.91 | -78.39 | 16.93 | 0.14 | 93.36 | 95.90 | ERR12713885 | This study |
| D5208 | *H. antiochus aranea* | Male | Venezuela | 7.68 | -72.23 | 12.08 | 0.16 | 92.23 | 95.49 | ERR12713886 | This study |
| D5221 | *H. antiochus aranea* | Male | Venezuela | 7.68 | -72.23 | 12.47 | 0.15 | 92.05 | 94.72 | ERR12713887 | This study |
| D5213 | *H. antiochus aranea* | Male | Venezuela | 7.68 | -72.23 | 15.56 | 0.15 | 92.00 | 95.60 | ERR12713888 | This study |
| **D5252** | ***H. antiochus araneides*** | **Male** | **Venezuela** | **8.11** | **-72.25** | **7.41** | **0.19** | **92.03** | **95.20** | ERR12713889 | This study |
| D5253 | *H. antiochus araneides* | Male | Venezuela | 8.11 | -72.25 | 13.24 | 0.16 | 91.67 | 95.52 | ERR12713890 | This study |
| D5259 | *H. antiochus araneides* | Male | Venezuela | 10.45 | -67.63 | 14.31 | 0.15 | 92.30 | 95.56 | ERR12713891 | This study |
| D5270 | *H. antiochus salvini* | Male | Venezuela | 7.31 | -61.47 | 13.43 | 0.15 | 91.73 | 95.34 | ERR12713892 | This study |
| D5273 | *H. antiochus salvini* | Female | Venezuela | 7.31 | -61.47 | 11.92 | 0.15 | 92.28 | 96.39 | ERR12713893 | This study |
| D5276 | *H. antiochus salvini* | Female | Venezuela | 7.31 | -61.47 | 13.03 | 0.15 | 92.02 | 95.40 | ERR12713894 | This study |
| A323 | *H. antiochus subsp nov.* | Female | Colombia | 5.62 | -72.30 | 17.03 | 0.14 | 89.63 | 95.57 | ERR12713895 | This study |
| R1163 | *H. antiochus subsp nov.* | Male | Colombia | 2.11 | -74.79 | 14.20 | 0.14 | 92.58 | 96.36 | ERR12713896 | This study |

| | | | | | | | | | | | |
|---|---|---|---|---|---|---|---|---|---|---|---|
| S4895 | *H. antiochus subsp nov.* | Female | Colombia | 2.58 | -72.71 | 22.07 | 0.13 | 93.16 | 95.74 | ERR12713897 | This study |
| A28 | *H. congener aquilionaris* | Male | Colombia | 1.18 | -76.66 | 14.05 | 0.15 | 90.42 | 94.60 | ERR12713898 | This study |
| A3740 | *H. congener aquilionaris* | Male | Colombia | 1.80 | -75.66 | 21.77 | 0.13 | 91.12 | 97.06 | ERR12713899 | This study |
| A3837 | *H. congener aquilionaris* | Male | Colombia | 3.50 | -74.02 | 18.67 | 0.14 | 91.20 | 97.09 | ERR12713900 | This study |
| A4278 | *H. congener aquilionaris* | Male | Colombia | 1.22 | -76.68 | 16.67 | 0.14 | 90.20 | 93.84 | ERR12713901 | This study |
| R806 | *H. congener congener* | Female | Ecuador | -1.44 | -78.42 | 13.59 | 0.15 | 91.83 | 97.74 | ERR12713902 | This study |
| R807 | *H. congener congener* | Male | Ecuador | -1.44 | -78.42 | 12.98 | 0.15 | 92.35 | 97.93 | ERR12713903 | This study |
| R808 | *H. congener congener* | Male | Ecuador | -1.44 | -78.42 | 12.48 | 0.15 | 92.16 | 97.69 | ERR12713904 | This study |
| R831 | *H. congener congener* | Female | Ecuador | -4.04 | -78.58 | 10.21 | 0.16 | 89.70 | 93.03 | ERR12713905 | This study |
| **R843** | ***H. congener congener*** | **Male** | **Ecuador** | **-4.14** | **-79.25** | **9.08** | **0.39** | **93.17** | **90.30** | ERR12713906 | This study |
| S40203 | *H. congener congener* | Male | Ecuador | -1.07 | -77.46 | 15.49 | 0.15 | 91.89 | 97.42 | ERR12713907 | This study |
| S857 | *H. eleuchia eleuchia* | Female | Panama | 8.86 | -79.88 | 17.05 | 0.15 | 93.30 | 97.36 | ERR12713911 | This study |
| S2083 | *H. eleuchia eleuchia* | Male | Colombia | 7.03 | -73.05 | 19.26 | 0.15 | 93.24 | 97.66 | ERR12713909 | This study |
| A2458 | *H. eleuchia eleuchia* | Male | Colombia | 5.08 | -74.57 | 13.74 | 0.16 | 90.79 | 96.29 | ERR12713910 | This study |
| S3106 | *H. eleuchia eleuchia* | Male | Colombia | 4.40 | -75.21 | 10.97 | 0.17 | 93.67 | 94.13 | ERR12713911 | This study |
| A3154 | *H. eleuchia eleuchia* | Male | Colombia | 3.68 | -76.53 | 17.98 | 0.15 | 90.23 | 96.68 | ERR12713912 | This study |
| S534 | *H. eleuchia elesinus* | Male | Colombia | 3.90 | -76.63 | 19.47 | 0.14 | 95.01 | 97.87 | ERR12713913 | This study |
| A3646 | *H. eleuchia elesinus* | Male | Colombia | 3.52 | -76.76 | 15.84 | 0.15 | 90.60 | 97.15 | ERR12713914 | This study |
| A4749 | *H. eleuchia elesinus* | Female | Colombia | 3.58 | -76.78 | 14.31 | 0.15 | 89.75 | 97.71 | ERR12713915 | This study |
| R4864 | *H. eleuchia elesinus* | Female | Colombia | 3.90 | 76.69 | 13.96 | 0.15 | 93.65 | 98.38 | ERR12713916 | This study |
| R4871 | *H. eleuchia elesinus* | Male | Colombia | 3.90 | 76.69 | 12.87 | 0.15 | 93.36 | 98.20 | ERR12713917 | This study |
| S8970 | *H. eleuchia elesinus* | Female | Panama | 7.55 | -78.20 | 15.22 | 0.14 | 92.72 | 94.75 | ERR12713918 | This study |
| S2852 | *H. eleuchia primularis* | Male | Ecuador | -2.44 | -79.05 | 14.71 | 0.15 | 92.91 | 97.68 | ERR12713919 | This study |
| R869 | *H. eleuchia primularis* | Male | Ecuador | -3.78 | -79.82 | 11.05 | 0.16 | 92.82 | 97.90 | ERR12713920 | This study |
| R870 | *H. eleuchia primularis* | Female | Ecuador | -3.78 | -79.82 | 12.02 | 0.15 | 92.53 | 97.69 | ERR12713921 | This study |
| R887 | *H. eleuchia primularis* | Female | Ecuador | -3.65 | -79.77 | 10.49 | 0.16 | 92.45 | 97.78 | ERR12713922 | This study |
| R888 | *H. eleuchia primularis* | Male | Ecuador | -3.65 | -79.79 | 11.06 | 0.16 | 92.53 | 97.95 | ERR12713923 | This study |
| S40343 | *H. eleuchia primularis* | Female | Ecuador | 0.08 | -78.89 | 14.57 | 0.15 | 92.85 | 97.83 | ERR12713924 | This study |
| D3 | *H. hewitsoni* | Male | Panama | 7.80 | -80.75 | 14.93 | 0.16 | 92.26 | 96.90 | ERR12713925 | This study |
| D6 | *H. hewitsoni* | Female | Panama | 7.80 | -80.75 | 12.05 | 0.16 | 92.40 | 98.04 | ERR12713926 | This study |
| D7 | *H. hewitsoni* | Male | Panama | 7.80 | -80.75 | 12.21 | 0.16 | 92.63 | 97.78 | ERR12713927 | This study |

| | | | | | | | | | | | |
|---|---|---|---|---|---|---|---|---|---|---|---|
| S33 | *H. leucadia pseudorhea* | | Peru | -10.68 | -75.12 | 17.00 | 0.10 | 94.12 | 97.11 | ERR12713929 | This study |
| A4016 | *H. leucadia pseudorhea* | Female | Colombia | -4.13 | -69.94 | 18.69 | 0.10 | 90.48 | 98.05 | ERR12713928 | This study |
| A2790 | *H. sapho sapho* | Male | Colombia | 6.85 | -73.03 | 17.77 | 0.13 | 90.22 | 91.86 | ERR12713930 | This study |
| R2794 | *H. sapho sapho* | Female | Colombia | 6.85 | -73.03 | 12.22 | 0.15 | 93.58 | 92.92 | ERR12713931 | This study |
| R4510 | *H. sapho sapho* | Female | Colombia | 7.06 | -73.38 | 15.25 | 0.14 | 94.14 | 98.22 | ERR12713932 | This study |
| R5091 | *H. sapho sapho* | Female | Colombia | 5.84 | 74.84 | 14.14 | 0.14 | 93.86 | 98.26 | ERR12713933 | This study |
| R5092 | *H. sapho sapho* | Female | Colombia | 5.84 | 74.84 | 14.57 | 0.13 | 92.87 | 97.98 | ERR12713934 | This study |
| R5237 | *H. sapho sapho* | Male | Panama | 9.12 | -79.49 | 12.96 | 0.14 | 92.91 | 91.76 | ERR12713935 | This study |
| R5314 | *H. sapho sapho* | Female | Panama | 9.12 | -79.49 | 12.89 | 0.14 | 92.55 | 97.81 | ERR12713936 | This study |
| R5324 | *H. sapho sapho* | Male | Panama | 9.12 | -79.49 | 13.87 | 0.14 | 92.09 | 97.71 | ERR12713937 | This study |
| A3497 | *H. sapho chocoensis* | Male | Colombia | 3.99 | -76.23 | 14.40 | 0.14 | 90.64 | 98.03 | ERR12713938 | This study |
| A3501 | *H. sapho chocoensis* | Male | Colombia | 3.99 | -76.23 | 17.35 | 0.14 | 90.55 | 97.42 | ERR12713939 | This study |
| R3510 | *H. sapho chocoensis* | Female | Colombia | 3.85 | -77.26 | 15.72 | 0.14 | 93.77 | 98.17 | ERR12713940 | This study |
| R3511 | *H. sapho chocoensis* | Male | Colombia | 3.85 | -77.26 | 13.07 | 0.15 | 93.60 | 98.42 | ERR12713941 | This study |
| A3491 | *H. sapho chocoensis* | Female | Colombia | 3.99 | -76.23 | 13.49 | 0.14 | 90.83 | 97.46 | ERR12713942 | This study |
| A4534 | *H. sara brevimaculata* | Female | Colombia | 3.94 | -77.36 | 24.55 | 0.04 | 90.63 | 96.14 | ERR12713943 | This study |
| A4538 | *H. sara brevimaculata* | Male | Colombia | 3.94 | -77.36 | 23.84 | 0.04 | 91.57 | 96.94 | ERR12713944 | This study |
| A4678 | *H. sara elektra* | Female | Colombia | 3.32 | -76.63 | 24.95 | 0.05 | 90.93 | 97.09 | ERR12713945 | This study |
| A4680 | *H. sara elektra* | Male | Colombia | 3.32 | -76.63 | 17.22 | 0.06 | 90.84 | 98.82 | ERR12713946 | This study |
| A4683 | *H. sara elektra* | Female | Colombia | 3.32 | -76.63 | 18.66 | 0.06 | 91.96 | 98.07 | ERR12713947 | This study |
| S4704 | *H. sara elektra* | Male | Colombia | 3.32 | -76.63 | 23.01 | 0.05 | 95.22 | 98.76 | ERR12713948 | This study |
| S29 | *H. sara magdalena* | Male | Panama | 8.71 | -79.91 | 19.60 | 0.03 | 94.31 | 98.68 | ERR12713949 | This study |
| R31 | *H. sara magdalena* | Male | Panama | 8.71 | -79.91 | 16.22 | 0.05 | 93.68 | 98.81 | ERR12713950 | This study |
| R32 | *H. sara magdalena* | Male | Panama | 8.71 | -79.91 | 16.79 | 0.05 | 93.15 | 98.69 | ERR12713951 | This study |
| S2172 | *H. sara magdalena* | Male | Venezuela | 7.80 | -72.20 | 25.42 | 0.05 | 95.25 | 98.86 | ERR12713952 | This study |
| S3533 | *H. sara magdalena* | Male | Colombia | 5.57 | -77.50 | 24.81 | 0.04 | 94.70 | 98.66 | ERR12713953 | This study |
| A3900 | *H. sara magdalena* | Male | Colombia | 6.37 | -77.38 | 20.49 | 0.04 | 91.86 | 98.66 | ERR12713954 | This study |
| R4134 | *H. sara magdalena* | Male | Colombia | 6.45 | -70.69 | 18.47 | 0.06 | 93.69 | 98.91 | ERR12713955 | This study |
| R4186 | *H. sara magdalena* | Male | Colombia | 5.74 | -74.23 | 14.59 | 0.05 | 94.31 | 99.03 | ERR12713956 | This study |
| R4187 | *H. sara magdalena* | Male | Colombia | 5.74 | -74.23 | 18.42 | 0.04 | 93.08 | 98.74 | ERR12713957 | This study |
| A4340 | *H. sara magdalena* | Male | Colombia | 6.10 | -67.49 | 17.67 | 0.06 | 91.87 | 98.34 | ERR12713958 | This study |

| | | | | | | | | | | | |
|---|---|---|---|---|---|---|---|---|---|---|---|
| S4346 | *H. sara magdalena* | Female | Colombia | 6.10 | -67.49 | 17.27 | 0.06 | 95.14 | 98.54 | ERR12713959 | This study |
| D5248 | *H. sara magdalena* | Male | Venezuela | 8.07 | -72.25 | 14.00 | 0.05 | 91.75 | 96.50 | ERR12713960 | This study |
| D5211 | *H. sara magdalena* | Male | Venezuela | 7.68 | -72.23 | 13.67 | 0.06 | 92.66 | 98.64 | ERR12713961 | This study |
| S1 | *H. sara sara* | Male | Brazil | -1.47 | -48.44 | 19.23 | 0.06 | 93.48 | 87.39 | ERR12713962 | This study |
| S5 | *H. sara sara* | Male | Peru | -10.65 | -75.11 | 15.96 | 0.06 | 94.41 | 77.38 | ERR12713963 | This study |
| S8 | *H. sara sara* | Male | Peru | -10.33 | -74.94 | 20.50 | 0.06 | 93.97 | 97.85 | ERR12713964 | This study |
| S12 | *H. sara sara* | Female | Brazil | -4.07 | -54.85 | 19.82 | 0.06 | 92.66 | 97.97 | ERR12713965 | This study |
| R15 | *H. sara sara* | Male | Brazil | -1.21 | -54.80 | 14.19 | 0.06 | 93.10 | 98.66 | ERR12713966 | This study |
| R16 | *H. sara sara* | Male | Brazil | -10.89 | -55.44 | 15.06 | 0.06 | 93.33 | 98.54 | ERR12713967 | This study |
| R35 | *H. sara sara* | Female | Peru | -5.29 | -80.68 | 14.33 | 0.06 | 93.95 | 98.46 | ERR12713968 | This study |
| R36 | *H. sara sara* | Female | Peru | -5.29 | -80.68 | 16.82 | 0.04 | 91.84 | 97.98 | ERR12713969 | This study |
| R37 | *H. sara sara* | Male | Peru | -5.29 | -80.68 | 14.74 | 0.05 | 92.87 | 97.74 | ERR12713970 | This study |
| S1053 | *H. sara sara* | Male | Colombia | 4.18 | -73.68 | 22.58 | 0.05 | 93.75 | 97.98 | ERR12713971 | This study |
| R812 | *H. sara sara* | Female | Ecuador | -3.91 | -79.74 | 14.35 | 0.06 | 92.79 | 98.67 | ERR12713972 | This study |
| R820 | *H. sara sara* | Male | Ecuador | -2.87 | -78.37 | 16.24 | 0.06 | 93.14 | 98.80 | ERR12713973 | This study |
| R828 | *H. sara sara* | Male | Ecuador | -4.04 | -78.58 | 15.14 | 0.06 | 92.79 | 98.73 | ERR12713974 | This study |
| R3467 | *H. sara sara* | Male | Colombia | 1.80 | -75.66 | 14.34 | 0.06 | 93.72 | 99.14 | ERR12713975 | This study |
| R3751 | *H. sara sara* | Male | Colombia | 1.80 | -75.66 | 15.77 | 0.06 | 93.41 | 98.77 | ERR12713976 | This study |
| A4121 | *H. sara sara* | Female | Colombia | -3.77 | -70.34 | 27.52 | 0.05 | 91.97 | 98.23 | ERR12713977 | This study |
| A4272 | *H. sara sara* | Female | Colombia | 1.21 | -76.68 | 18.39 | 0.06 | 92.39 | 98.06 | ERR12713978 | This study |
| A4343 | *H. sara sara* | Female | Colombia | 6.10 | -67.49 | 19.93 | 0.06 | 91.63 | 98.30 | ERR12713979 | This study |
| A4489 | *H. sara sara* | Male | Ecuador | -1.04 | -77.84 | 25.07 | 0.05 | 92.34 | 97.96 | ERR12713980 | This study |
| S21204 | *H. sara sara* | Male | Surinam | 4.74 | -56.80 | 20.55 | 0.06 | 94.61 | 98.41 | ERR12713981 | This study |
| S40219 | *H. sara sara* | Male | Ecuador | -1.04 | -77.84 | 20.75 | 0.06 | 93.64 | 98.21 | ERR12713982 | This study |
| R871 | *H. sara sprucei* | Male | Ecuador | -3.77 | -79.83 | 15.05 | 0.05 | 92.73 | 97.60 | ERR12713983 | This study |
| R886 | *H. sara sprucei* | Male | Ecuador | -3.65 | -79.76 | 16.13 | 0.05 | 92.66 | 98.84 | ERR12713984 | This study |
| A3856 | *H. sara sprucei* | Female | Colombia | 2.96 | -78.18 | 23.72 | 0.03 | 91.74 | 98.77 | ERR12713985 | This study |
| A3858 | *H. sara sprucei* | Female | Colombia | 2.97 | -78.18 | 23.52 | 0.04 | 92.11 | 98.71 | ERR12713986 | This study |
| A3861 | *H. sara sprucei* | Male | Colombia | 2.96 | -78.18 | 22.83 | 0.04 | 91.44 | 98.70 | ERR12713987 | This study |
| R4590 | *H. sara sprucei* | Female | Colombia | 3.94 | -77.36 | 15.56 | 0.05 | 93.09 | 98.70 | ERR12713988 | This study |
| R4591 | *H. sara sprucei* | Male | Colombia | 3.94 | -77.36 | 13.71 | 0.05 | 93.31 | 98.84 | ERR12713989 | This study |

| | | | | | | | | | | | |
|---|---|---|---|---|---|---|---|---|---|---|---|
| R4597 | *H. sara sprucei* | Female | Colombia | 3.94 | -77.36 | 15.60 | 0.05 | 93.56 | 98.73 | ERR12713990 | This study |
| SRR4032025 | *H. charitonia* | | Puerto Rico | 18.02 | -66.83 | 16.57 | 0.06 | 91.60 | 79.23 | SAMN05224120 | Van Belleghem et al. 2017 |
| SRR4032026 | *H. charitonia* | | Puerto Rico | 18.02 | -66.83 | 14.35 | 0.06 | 90.53 | 80.29 | SAMN05224121 | Van Belleghem et al. 2017 |
| SAMEA8947140* | *H. sara* | Female | Panama | 9.12 | -79.70 | | | | | SAMEA8947140 | This study |
| SAMEA8947139* | *H. sara* | Female | Panama | 9.12 | -79.70 | | | | | SAMEA8947139 | This study |
| SAMEA112329098** | *H. congener* | Female | Colombia | 1.20 | -76.68 | | | | | SAMEA112329098 | This study |
| SAMEA112696452** | *H. sapho* | Male | Ecuador | 0.11 | -79.12 | | | | | SAMEA112696452 | This study |

*These specimens were used to generate the reference genome of *Heliconius sara*.

**These specimens were used to generate the HiC data.

**Table S2.** Lepidoptera genome assembly statistics modified from (1). We added our new *H. sara* assembly and the *H. charithonia* assembly from (2).

| | Scaffolds | Total length | Mean scaffold size | Maximum scaffold size | Scaffold N50 | Scaffold L50 | Contings | Contig N50 |
|---|---|---|---|---|---|---|---|---|
| ***Heliconius sara*** | 384 | 348,900,000 | 16,148,070 | 22,417,933 | 17,800,000 | 9 | 521 | 8,200,000 |
| *Heliconius charithonia* | 224 | 356,200,000 | 15,588,749 | 34,286,314 | 17,100,000 | 9 | *359* | 16,400,000 |
| *Heliconius erato* | 196 | 382,844,248 | 1,953,287 | 23,854,410 | 10,745,484 | 12 | *5,851* | 169,806 |
| *Heliconius melpomene* v2.5 | 332 | 275,198,613 | 18,000,000 | 18,000,000 | 14,300,000 | 34 | 3,126 | 328,886 |
| *Bombyx mori* | 43,462 | 481,803,763 | 11,085 | 16,203,812 | 4,008,358 | 38 | 87,972 | 15,765 |
| *Danaus plexippus* | 5,397 | 248,564,116 | 46,055 | 6,243,218 | 715,606 | 101 | 10,545 | 113,903 |
| *Lerema accius* | 29,988 | 298,173,436 | 9,943 | 3,082,282 | 525,349 | 160 | 52,985 | 18,018 |
| *Melitaea cinxia* | 8,261 | 389,907,520 | 47,198 | 668,473 | 119,328 | 970 | 45,618 | 15,003 |
| *Papilio glaucus* | 68,029 | 375,987,417 | 5,526 | 1,977,235 | 230,299 | 421 | 96,532 | 12,958 |
| *Papilio polytes* | 3,873 | 227,005,758 | 58,612 | 9,881,032 | 3,672,263 | 21 | 13,441 | 51,561 |
| *Papilio xuthus* | 5,572 | 243,890,167 | 43,770 | 16,292,344 | 6,198,915 | 16 | 10,483 | 133,779 |
| *Plutella xylostella* | 1,819 | 394,062,517 | 216,636 | 3,493,687 | 737,182 | 155 | 15,764 | 59,184 |
| *Bicyclus anynana* | 15,180 | 458,610,584 | 30,211 | 2,943,548 | 364,913 | 303 | 28,866 | 60,000 |
| *Chilo suppressalis* | 80,479 | 372,375,373 | 4,626 | 111,673 | 5,215 | 19,910 | 331,320 | 2,183 |
| *Manduca sexta* | 20,871 | 419,424,771 | 20,096 | 3,253,989 | 664,006 | 169 | 35,212 | 51,909 |
| *Plodia interpunctella* | 10,542 | 381,952,380 | 36,231 | 7,207,896 | 1,270,674 | 76 | 17,231 | 338,910 |

**Table S3.** BUSCO results statistics modified from (1). We added our *H. sara* assembly and the *H. charithonia* assembly of (2).

| | Complete Single-Copy BUSCOs % | Complete Duplicated BUSCOs % | Fragmented BUSCOs % | Missing BUSCOs % |
|---|---|---|---|---|
| ***Heliconius sara*** | 98 | 0.2 | 0.7 | 0.9 |
| *Heliconius charithonia* | 99 | 3.3 | 0.2 | 1.1 |
| *Heliconius erato* | 82 | 2.3 | 11 | 6.3 |
| *Heliconius melpomene* v1 | 81 | 2.8 | 11 | 7.2 |
| *Heliconius melpomene* v2.5 | 98 | 1.3 | 0.3 | 5 |
| *Bombyx mori* | 75 | 2.2 | 16 | 8.4 |
| *Danaus plexippus* | 87 | 3.5 | 10 | 2.7 |
| *Lerema accius* | 77 | 2.6 | 13 | 8.3 |
| *Melitaea cinxia* | 55 | 1.6 | 20 | 23 |
| *Papilio glaucus* | 75 | 2.7 | 14 | 9.6 |
| *Papilio polytes* | 76 | 2.4 | 12 | 11 |
| *Papilio xuthus* | 84 | 3 | 8.2 | 7.4 |
| *Plutella xylostella* | 74 | 20 | 11 | 13 |
| *Bicyclus anynana* | 81 | 3 | 12 | 6.5 |
| *Chilo suppressalis* | 33 | 0.7 | 17 | 48 |
| *Manduca sexta* | 81 | 4.4 | 11 | 6.4 |
| *Plodia interpunctella* | 85 | 3.4 | 9.4 | 4.7 |

### Figures

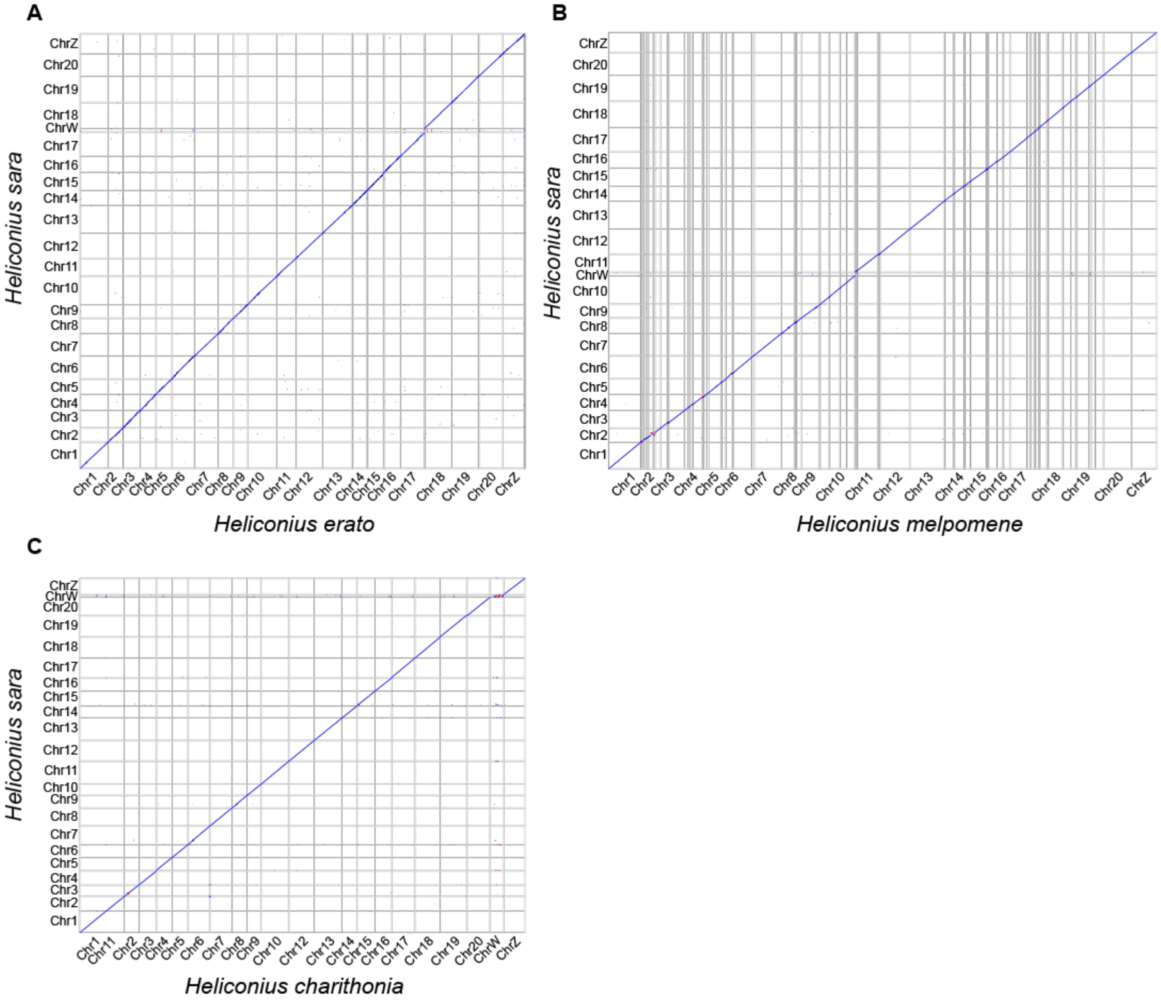


**Fig S1. Synteny plots showing high collinearity between *Heliconius* genomes*.*** Pairwise alignment between chromosomes of (A) *H. sara* and *H. melpomene*, (B) *H. sara* and *H. erato*, and (C) *H. sara* and *H. charithonia*. The GenBank accession numbers for the genomes of *H. melpomene*, *H. erato* and *H. charithonia* are GCA_000313835.2, GCA_018249695.1 and GCA_030704555.1, respectively. The W chromosome in *H. sara* corresponds to a single homolog in H. *charithonia*. Because the W chromosome was not assembled in the genomes of *H. erato* and *H. melpomene* genomes, we could not compare the W of *H. sara* W against them.

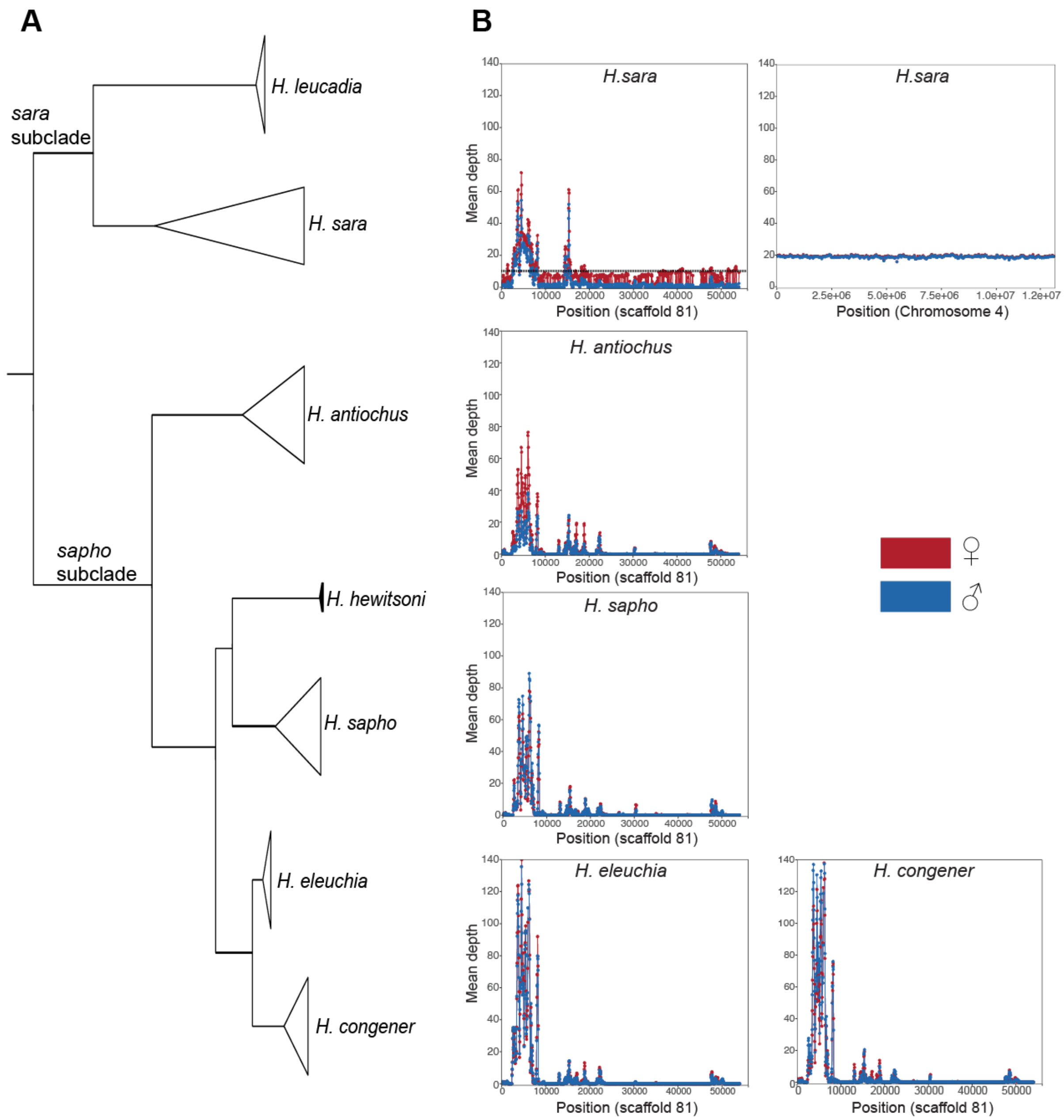


**Fig S2. Identification of the W chromosome in the genome of *H. sara*.** (A) Genome-wide topology of the *sara*-*sapho* clade. (B) Mean depth vs. position along the scaffold in each species. We plotted one (scaffold 81) out of the 32 scaffolds where females of the species *H. sara* showed half the sequencing depth of autosomes and where males do not map. The mean depth of an autosomal chromosome (Chr4) of the species *H. sara* is provided as an example for comparison with scaffold 81. Females are shown in red and males in blue.

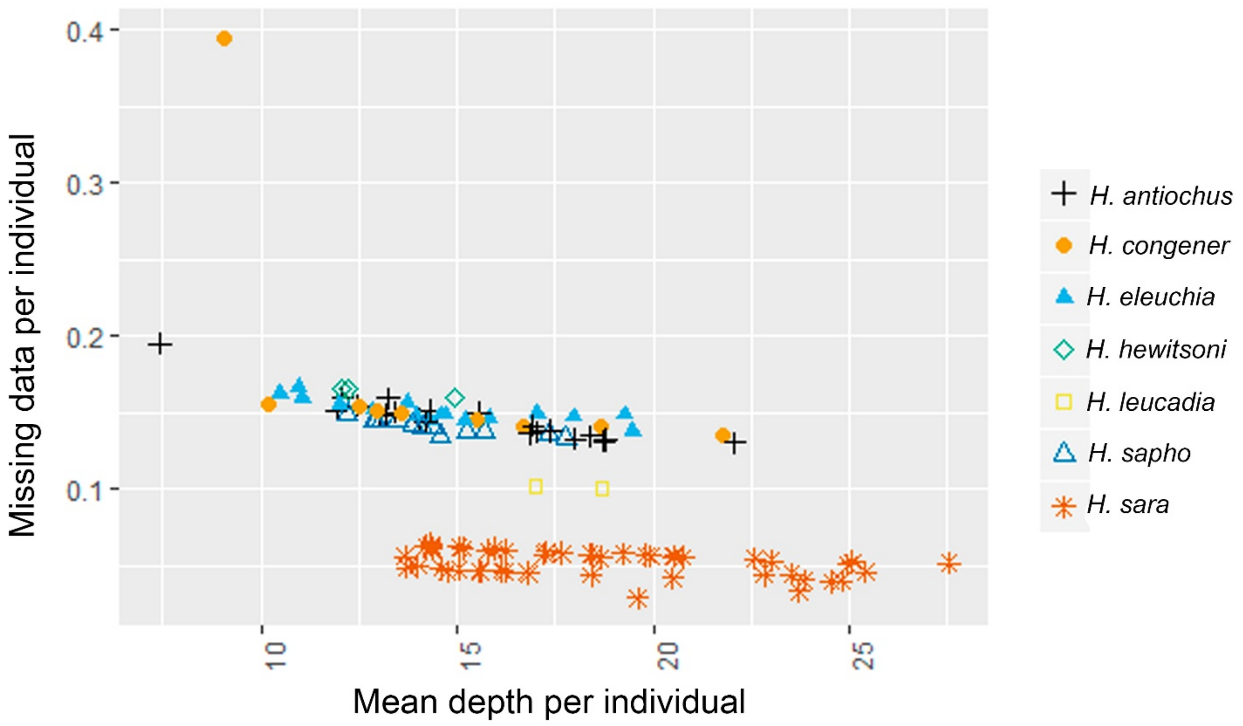


**Fig S3. Missing data and mean depth per individual.** Each species is symbolised by a unique symbol and colour. Note that the lower missing data proportion in *H. sara* is likely due to its similarity with the reference genome (*H. sara* female).

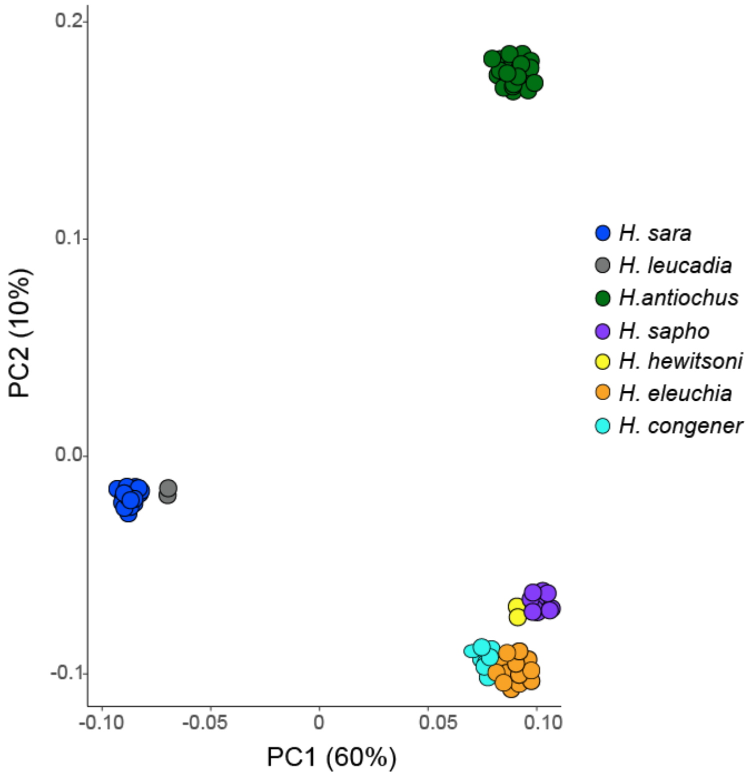


**Fig S4. Principal Component Analysis (PCA), performed with 3,685,916 SNPs.** The PCA groups the individuals into two main groups: (i) *H. sara and leucadia* (hereafter *sara* subclade), and (ii) *H. antiochus*, *H. eleuchia, H. congener, H. sapho* and *H. hewitsoni* (hereafter *sapho* subclade). The first two principal components explain 60% (PC1) and 10% (PC2) of the total variance, respectively. PC1 separates the *sara* subclade from the *sapho* subclade, whereas PC2 separates *H. antiochus* from the rest of the species of the *sapho* subclade. *H. sapho* is closer to *H. hewitsoni,* whereas *H. eleuchia* is closer to *H. congener.*

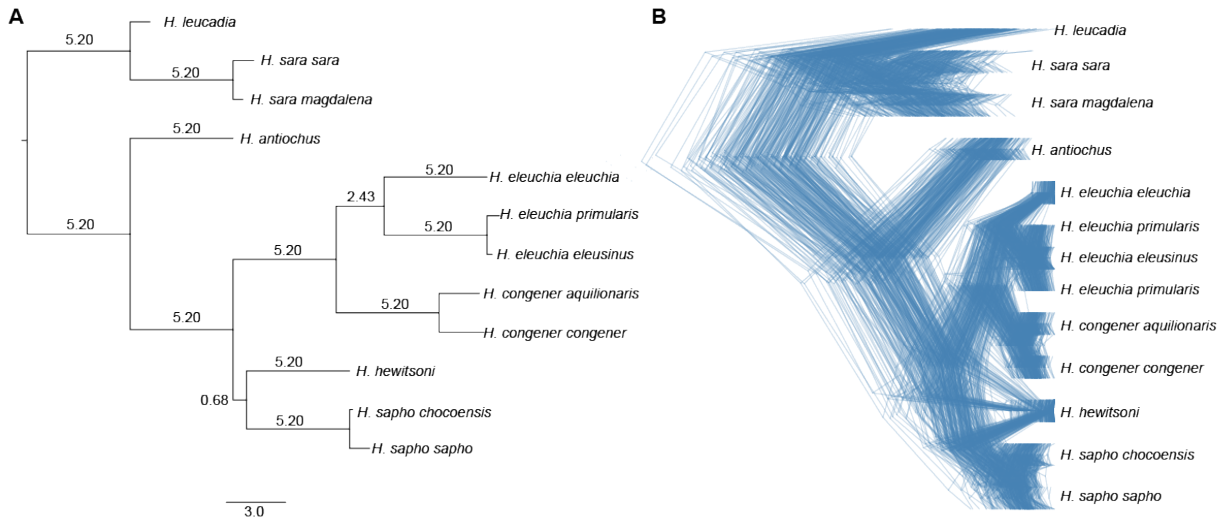


**Fig S5. Species tree based on ASTRAL multi-species coalescence and Densitree**. (A) ASTRAL species tree based on 271 phylogenetic trees (each recovered from a block of 100 loci). Branch lengths are shown in coalescent units. All nodes are supported with a bootstrap value of 100%. (B) DensiTree calculated from 271 topologies showing phylogenetic discordance within the *sara-sapho* clade.

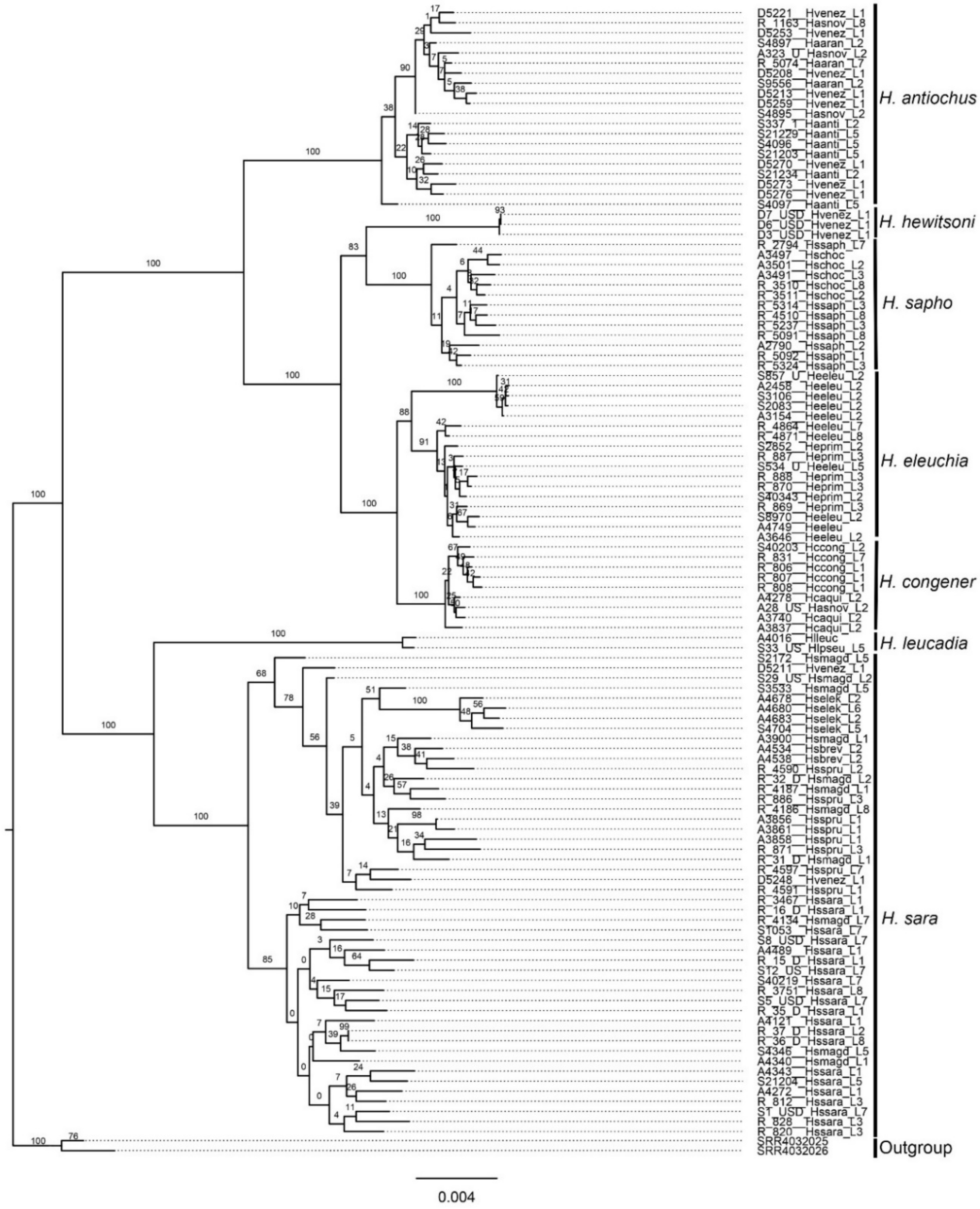


**Fig S6. Maximum likelihood phylogeny of chromosome 1.** Bootstrap support values are indicated at branches.

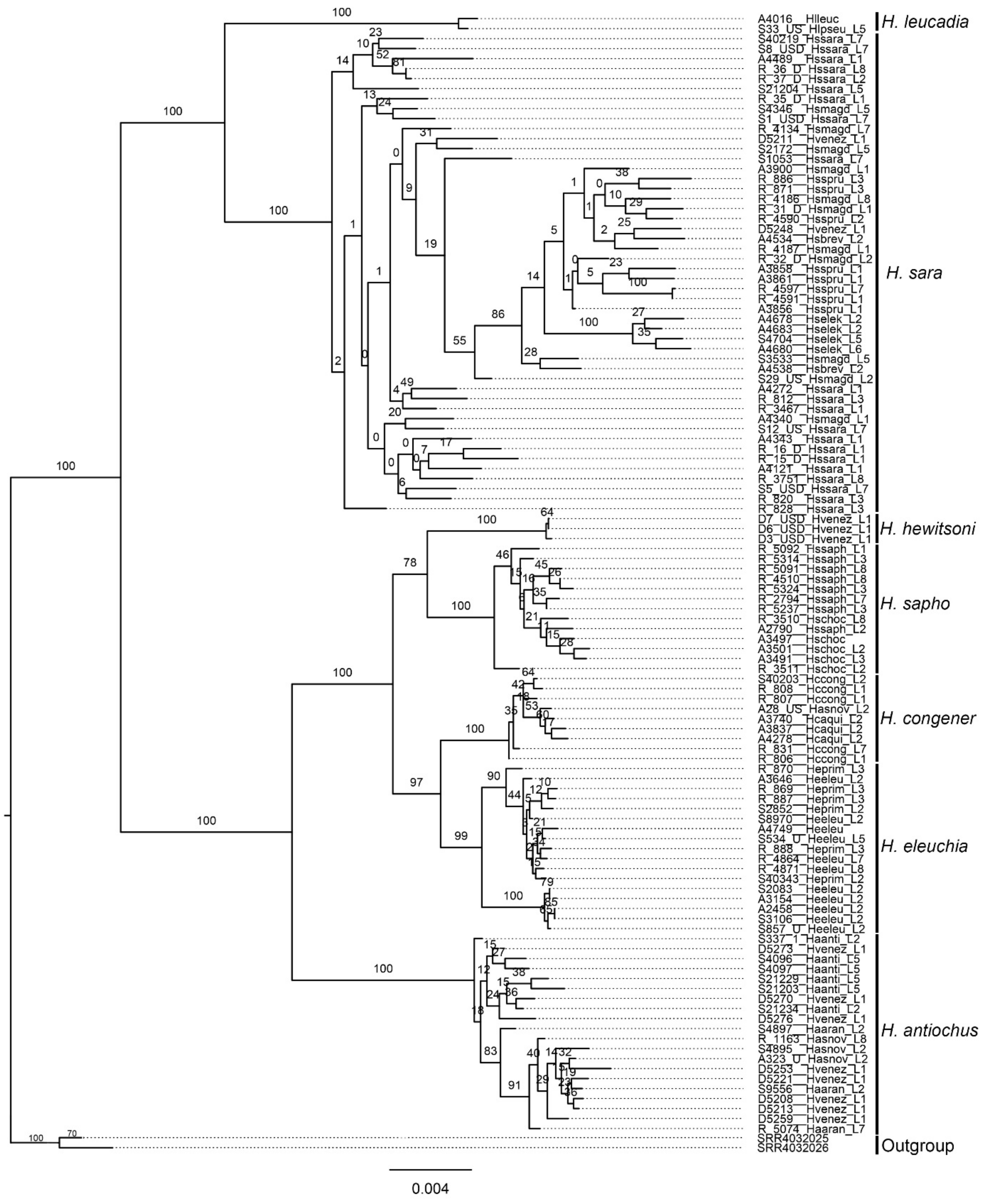


**Fig S7. Maximum likelihood phylogeny of chromosome 2.** Bootstrap support values are indicated at branches.

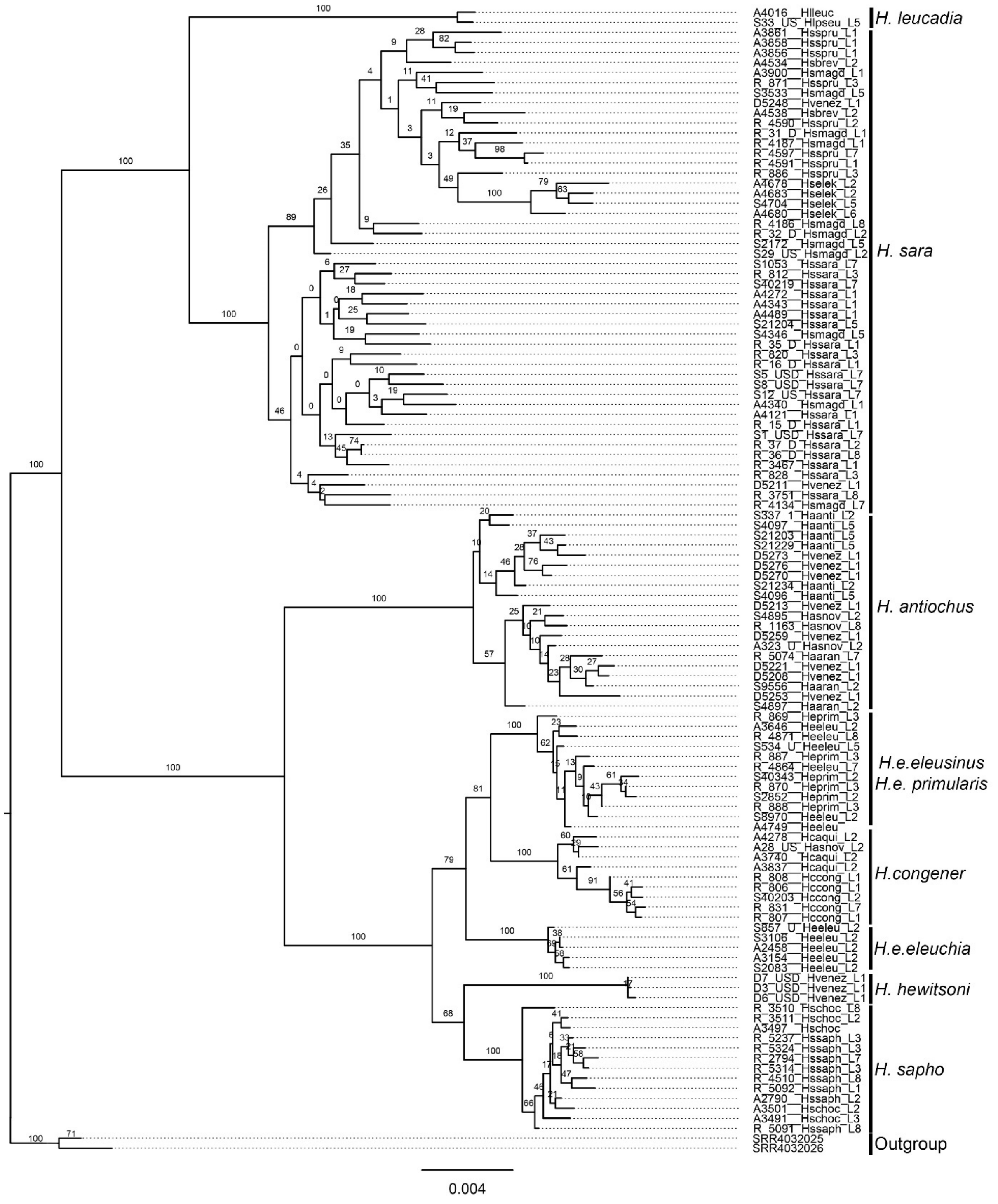


**Fig S8. Maximum likelihood phylogeny of chromosome 3.** Bootstrap support values are indicated at branches.

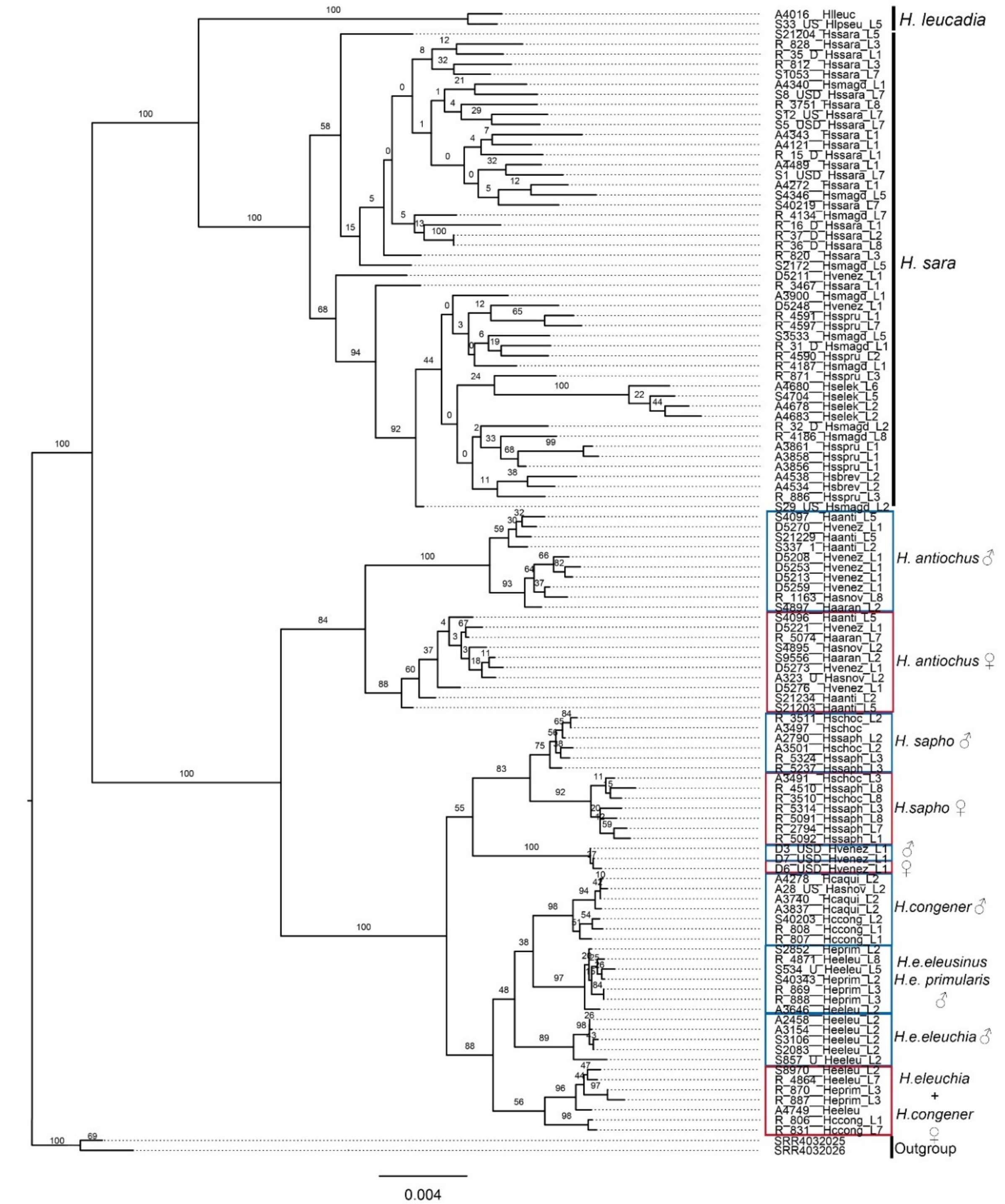


**Fig S9. Maximum likelihood phylogeny of chromosome 4.** Bootstrap support values are indicated at branches. Blue squares group males and while red squares group females.

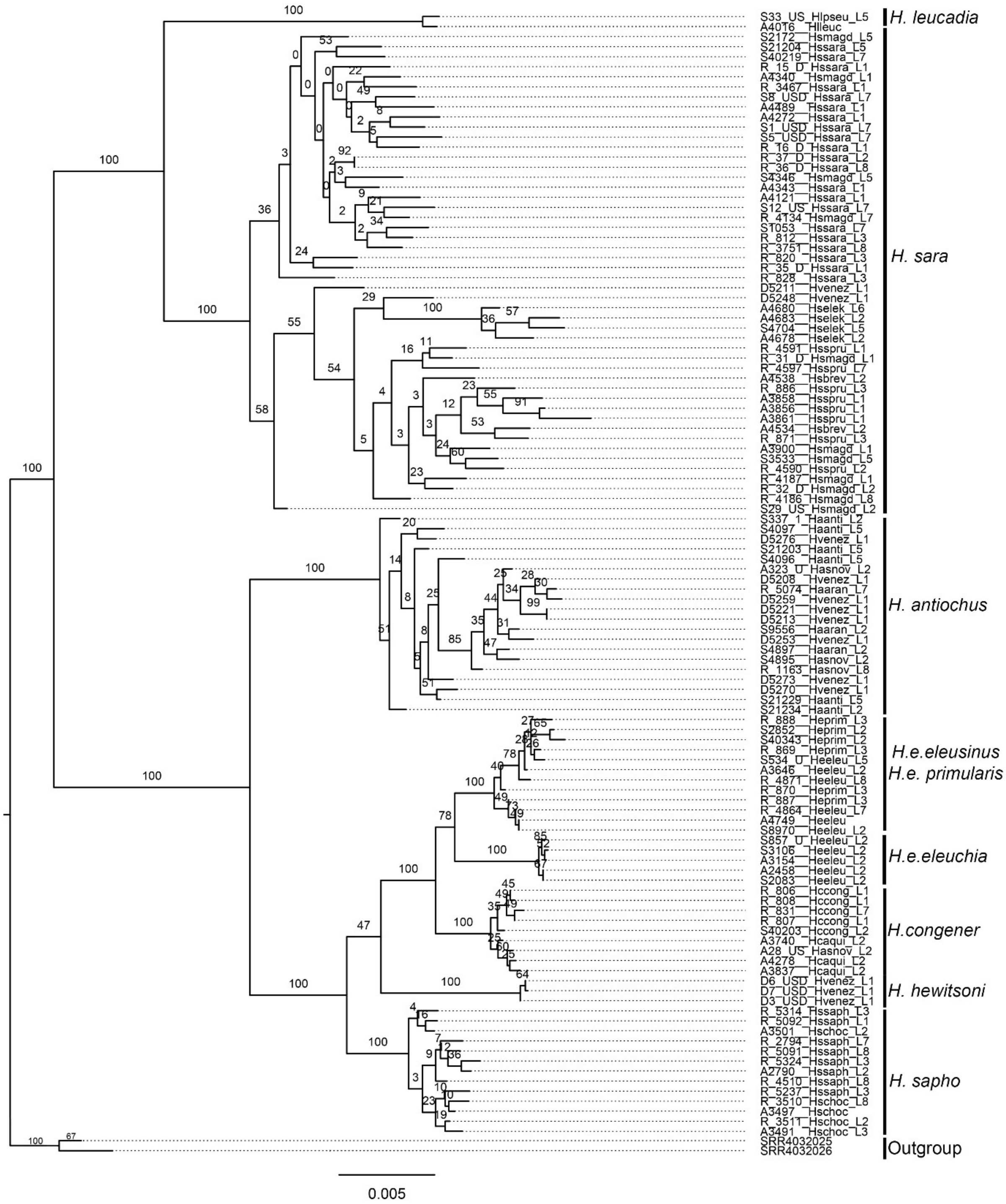


**Fig S10. Maximum likelihood phylogeny of chromosome 5.** Bootstrap support values are indicated at branches.

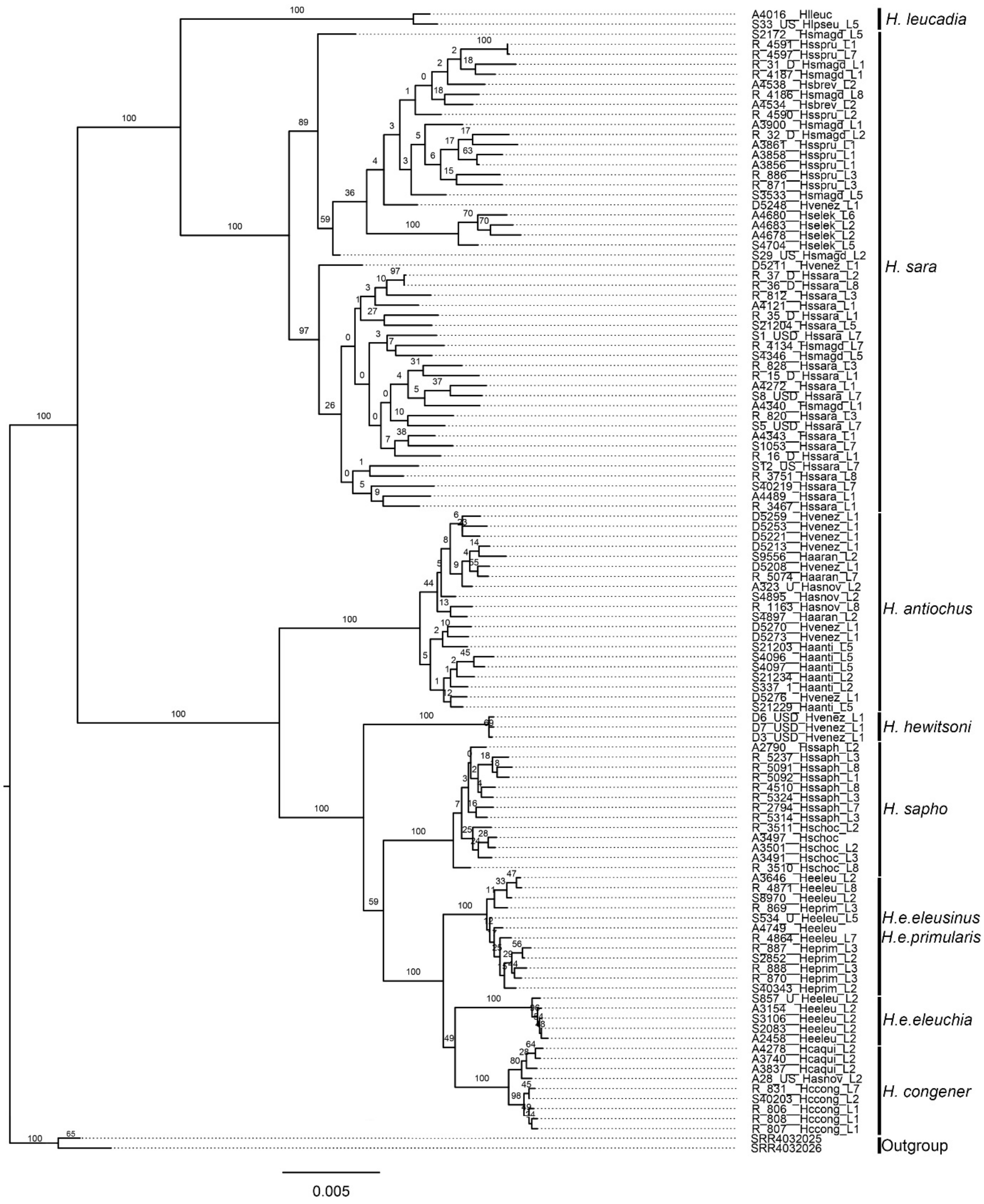


**Fig S11. Maximum likelihood phylogeny of chromosome 6.** Bootstrap support values are indicated at branches.

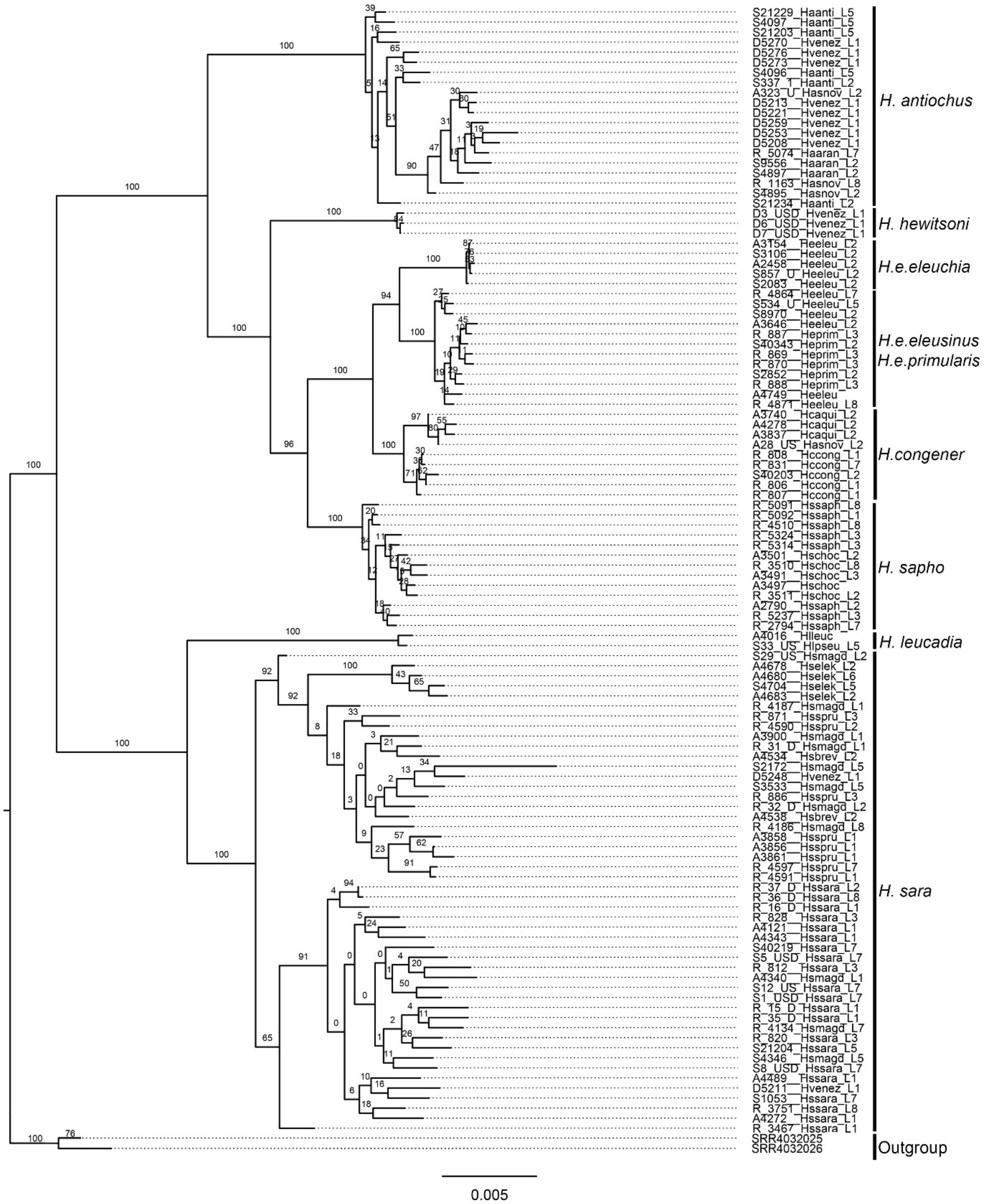


**Fig S12. Maximum likelihood phylogeny of chromosome 7.** Bootstrap support values are indicated at branches.

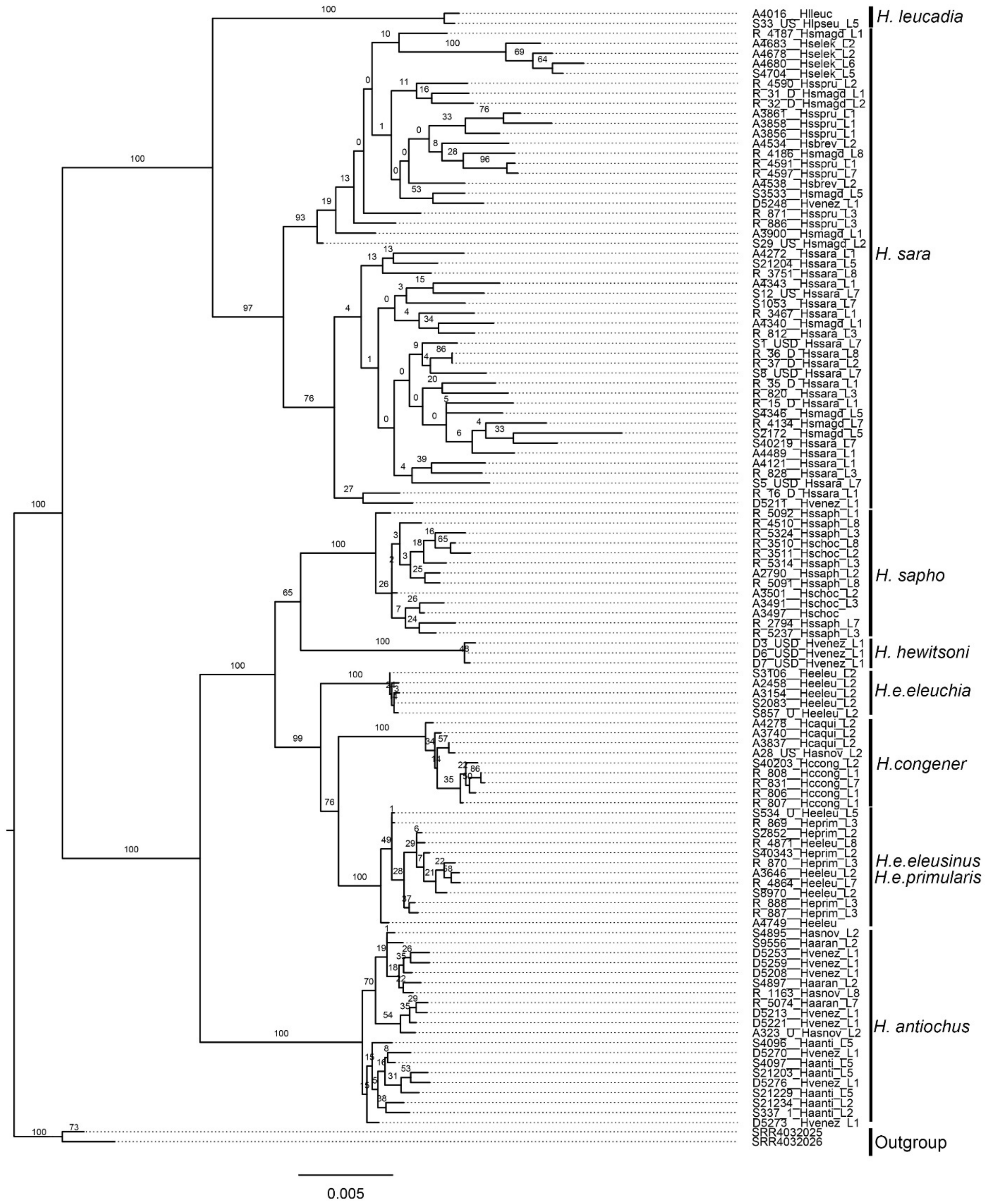


**Fig S13. Maximum likelihood phylogeny of chromosome 8.** Bootstrap support values are indicated at branches.

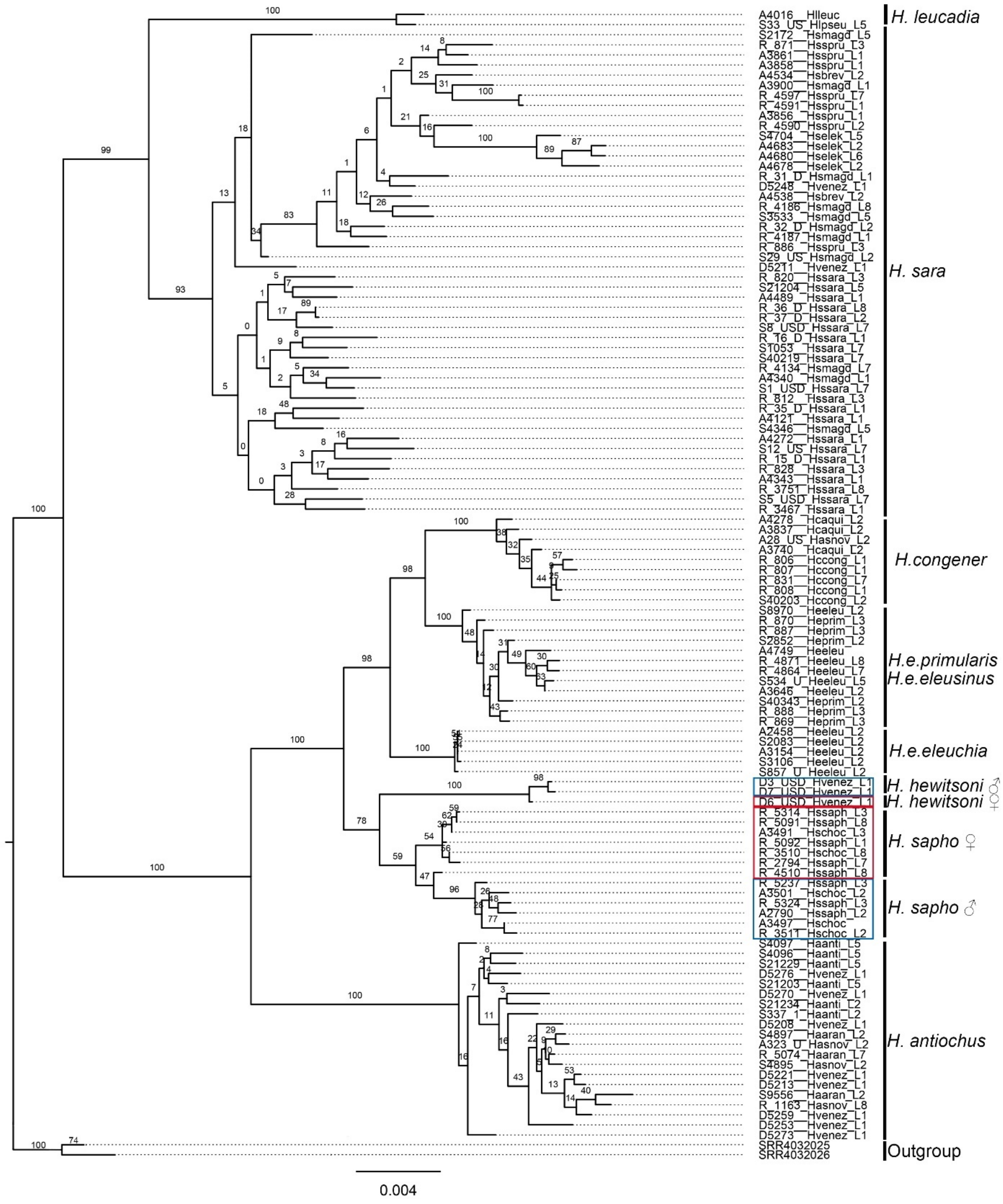


**Fig S14. Maximum likelihood phylogeny of chromosome 9.** Bootstrap support values are indicated at branches. Blue squares group males and while red squares group females.

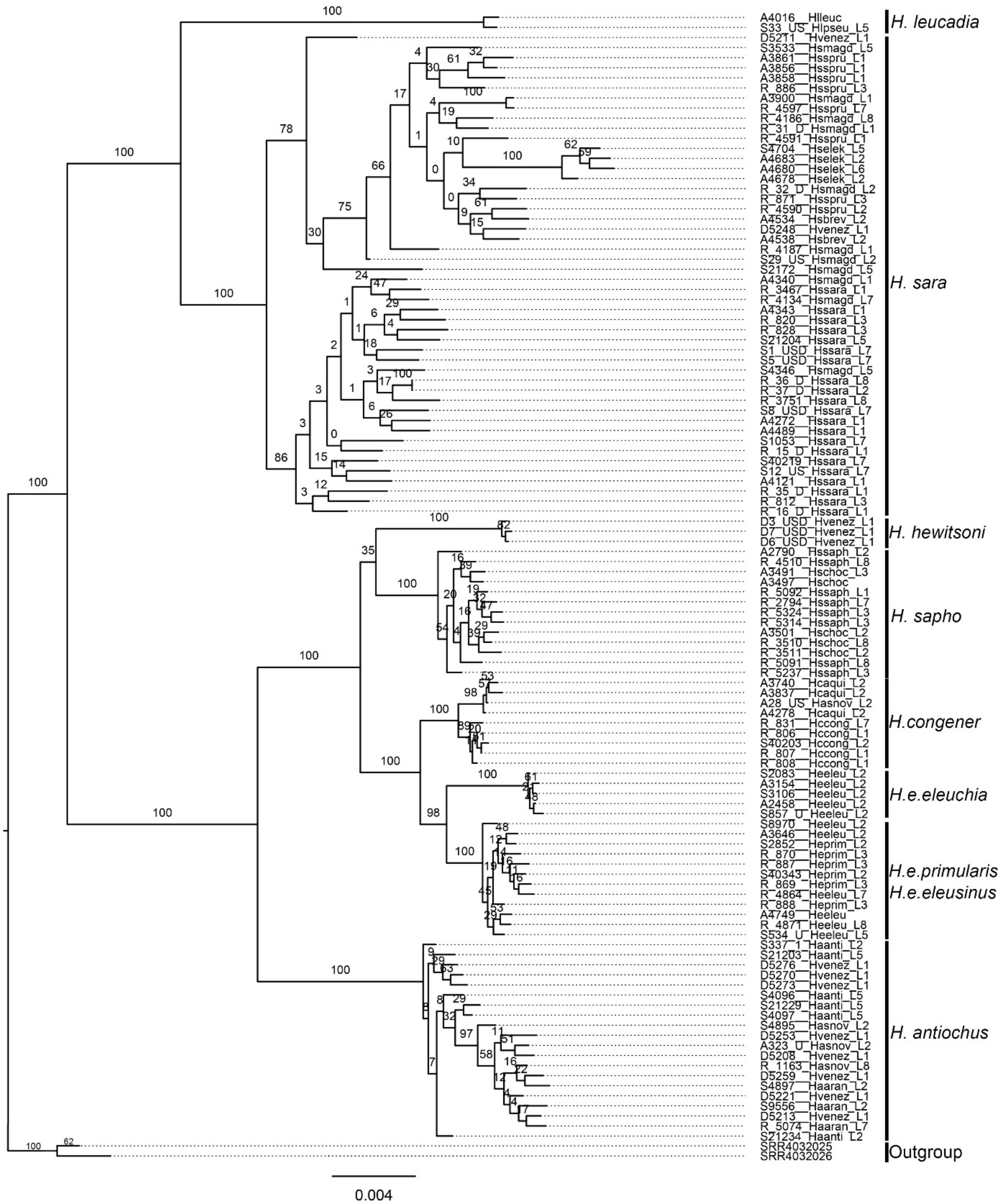

**Fig S15. Maximum likelihood phylogeny of chromosome 10.** Bootstrap support values are indicated at branches.

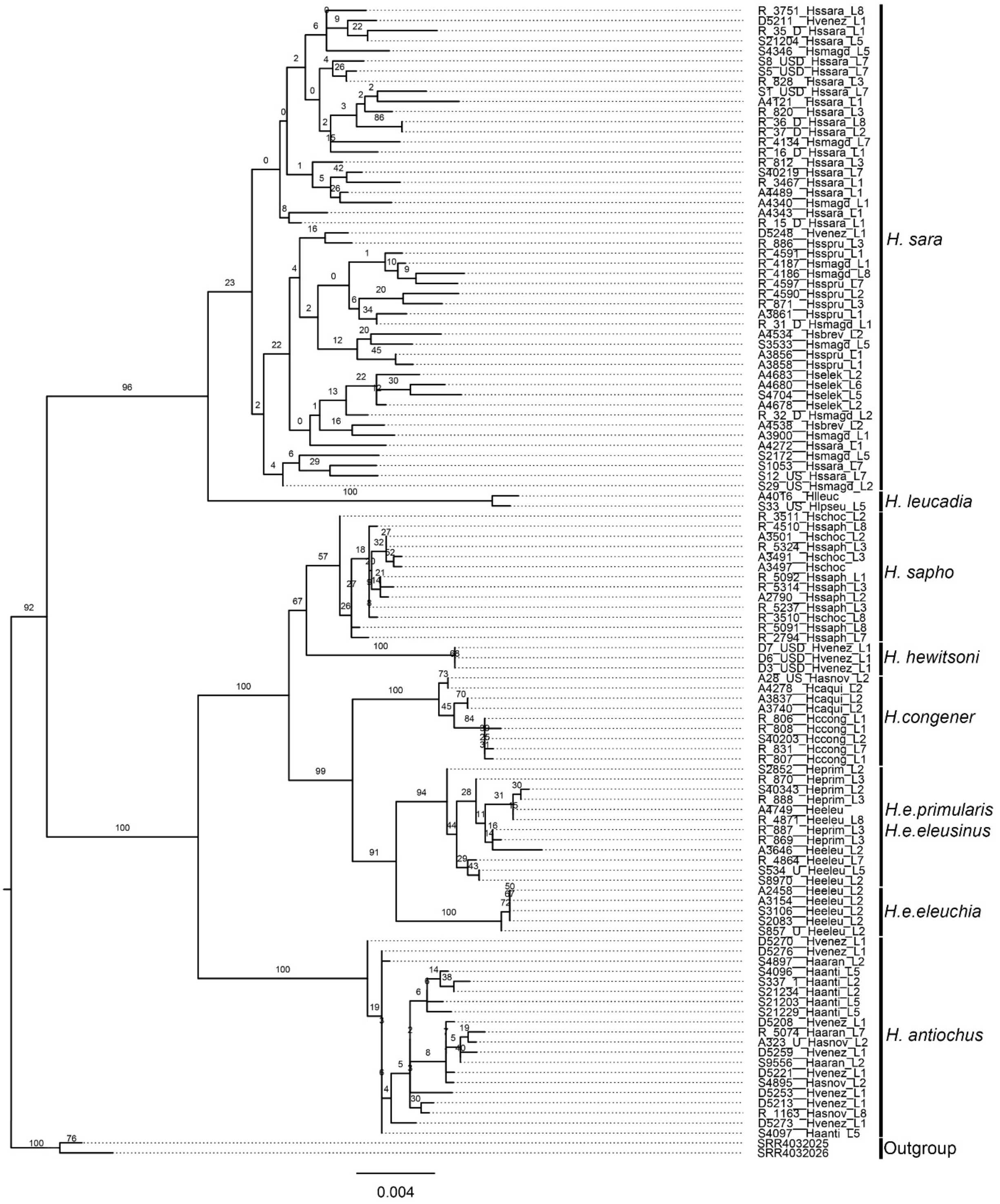


**Fig S16. Maximum likelihood phylogeny of chromosome 11.** Bootstrap support values are indicated at branches.

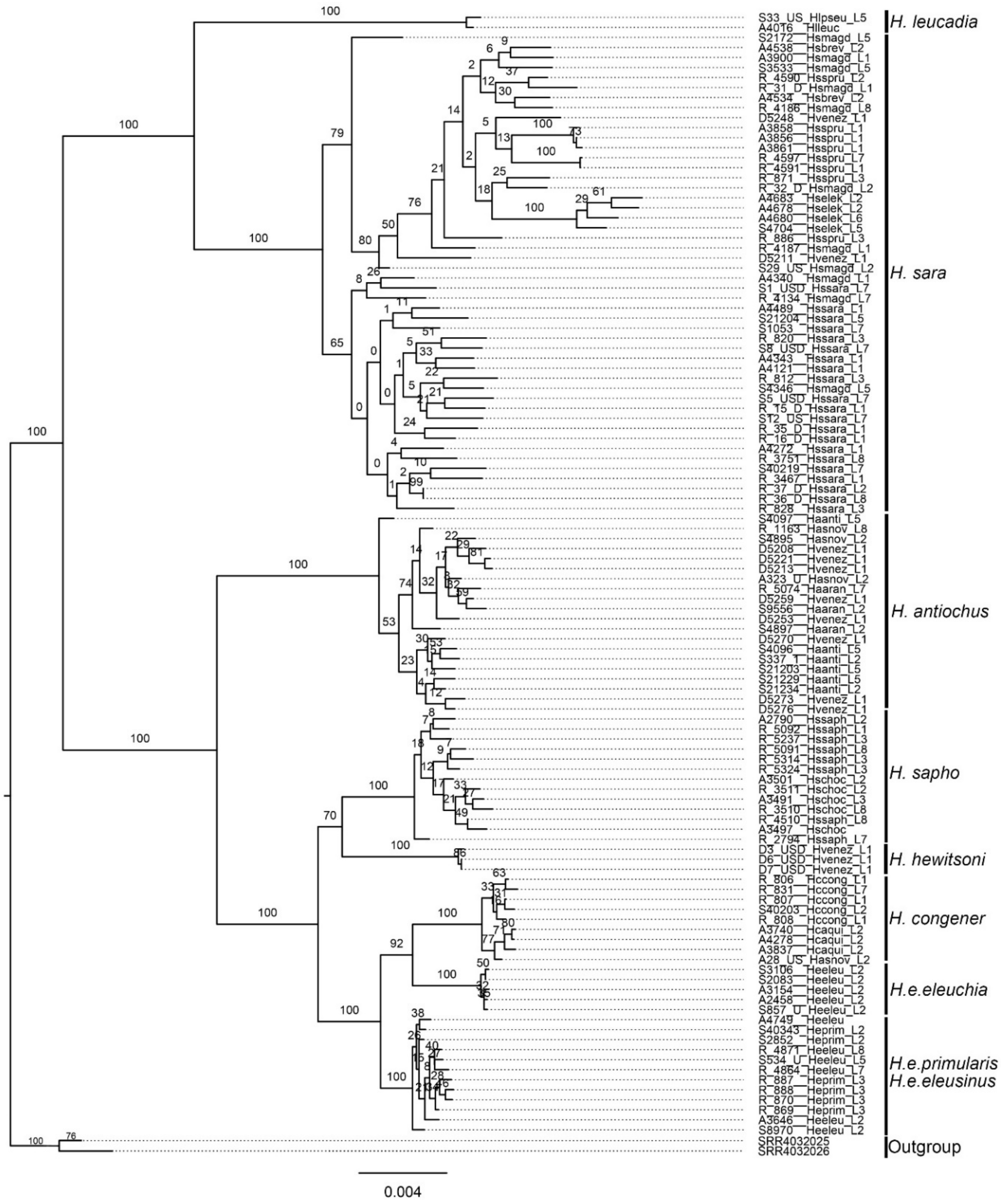


**Fig S17. Maximum likelihood phylogeny of chromosome 12.** Bootstrap support values are indicated at branches.

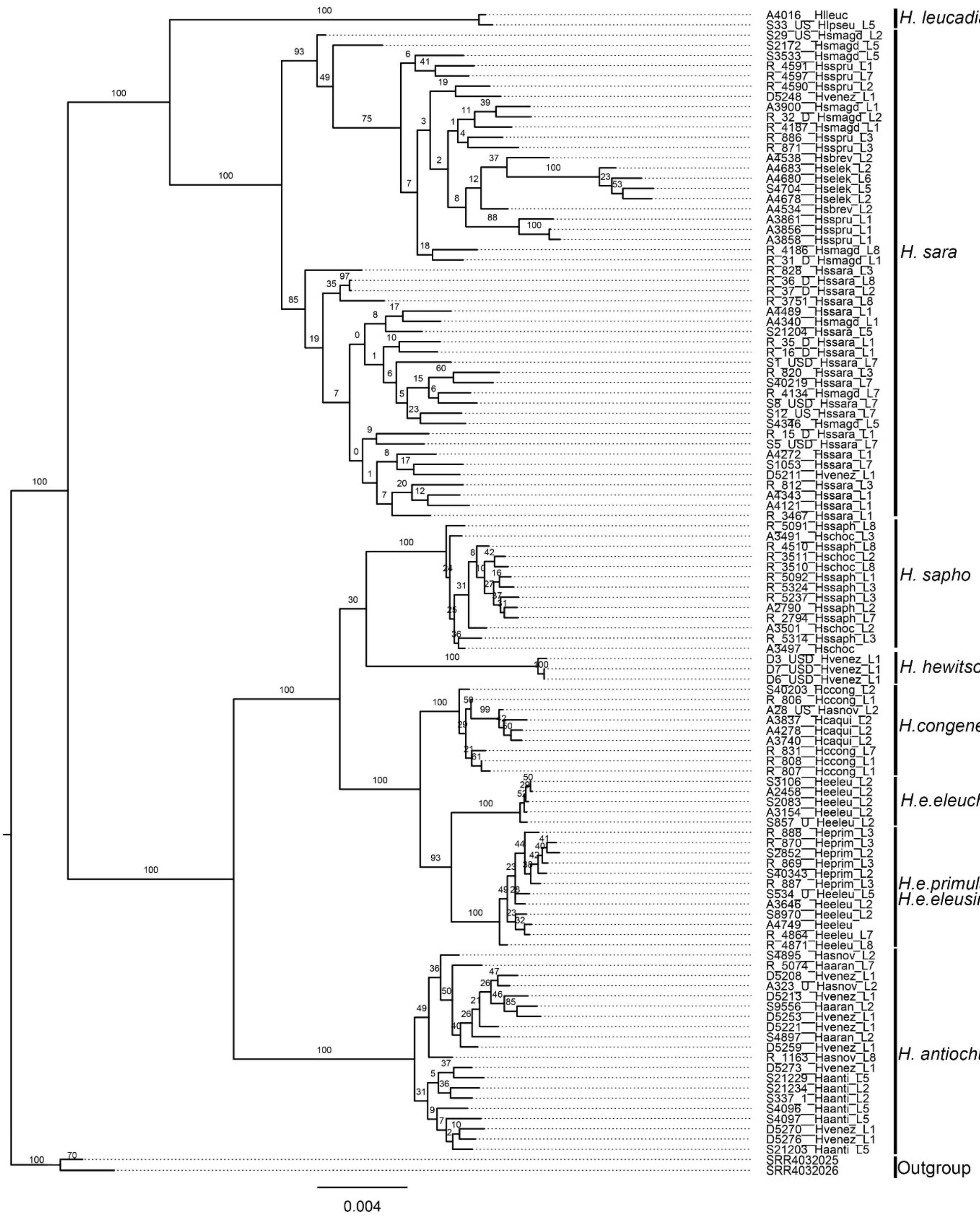


**Fig S18. Maximum likelihood phylogeny of chromosome 13.** Bootstrap support values are indicated at branches.

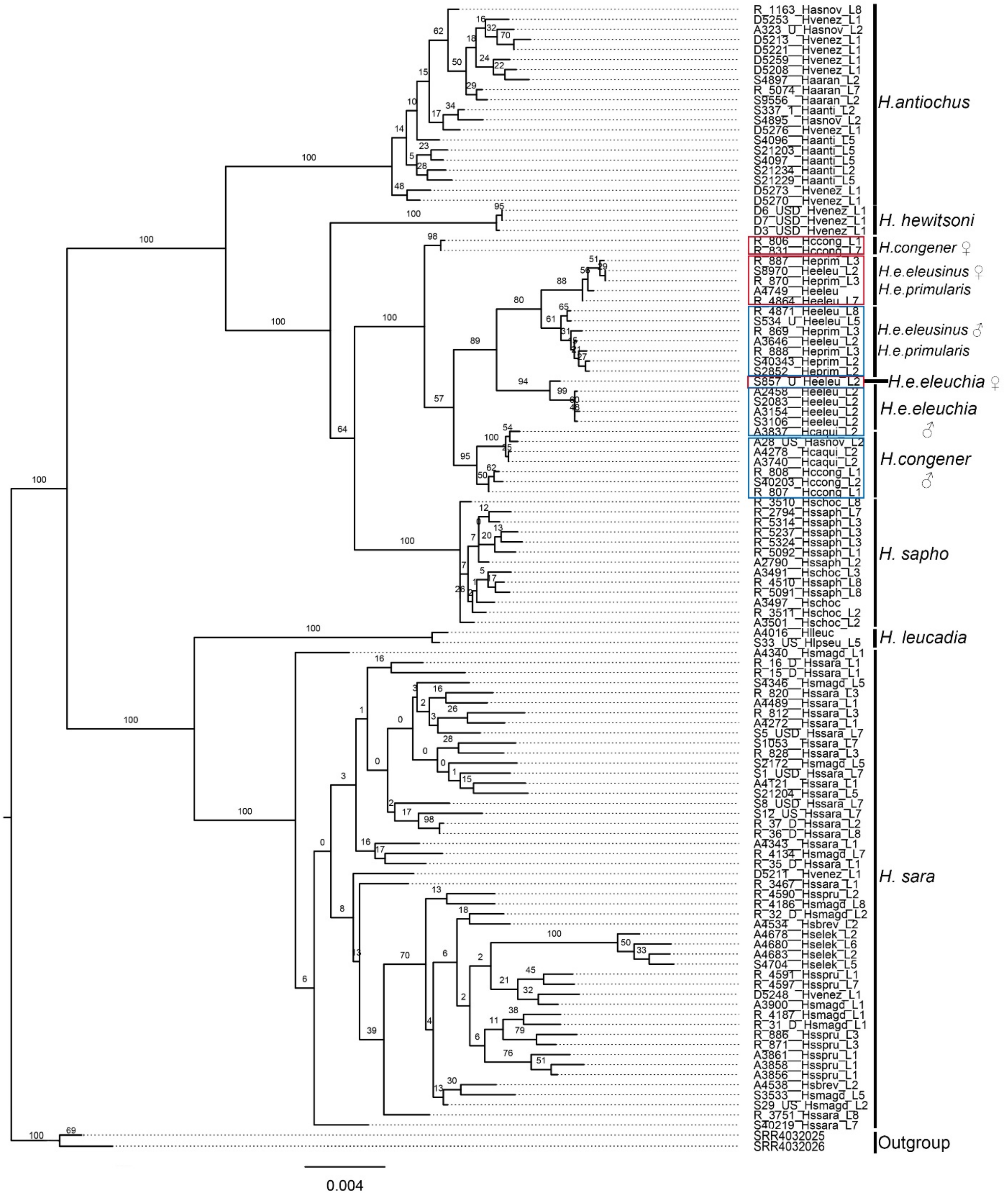


**Fig S19. Maximum likelihood phylogeny of chromosome 14.** Bootstrap support values are indicated at branches. Blue squares group males and while red squares group females.

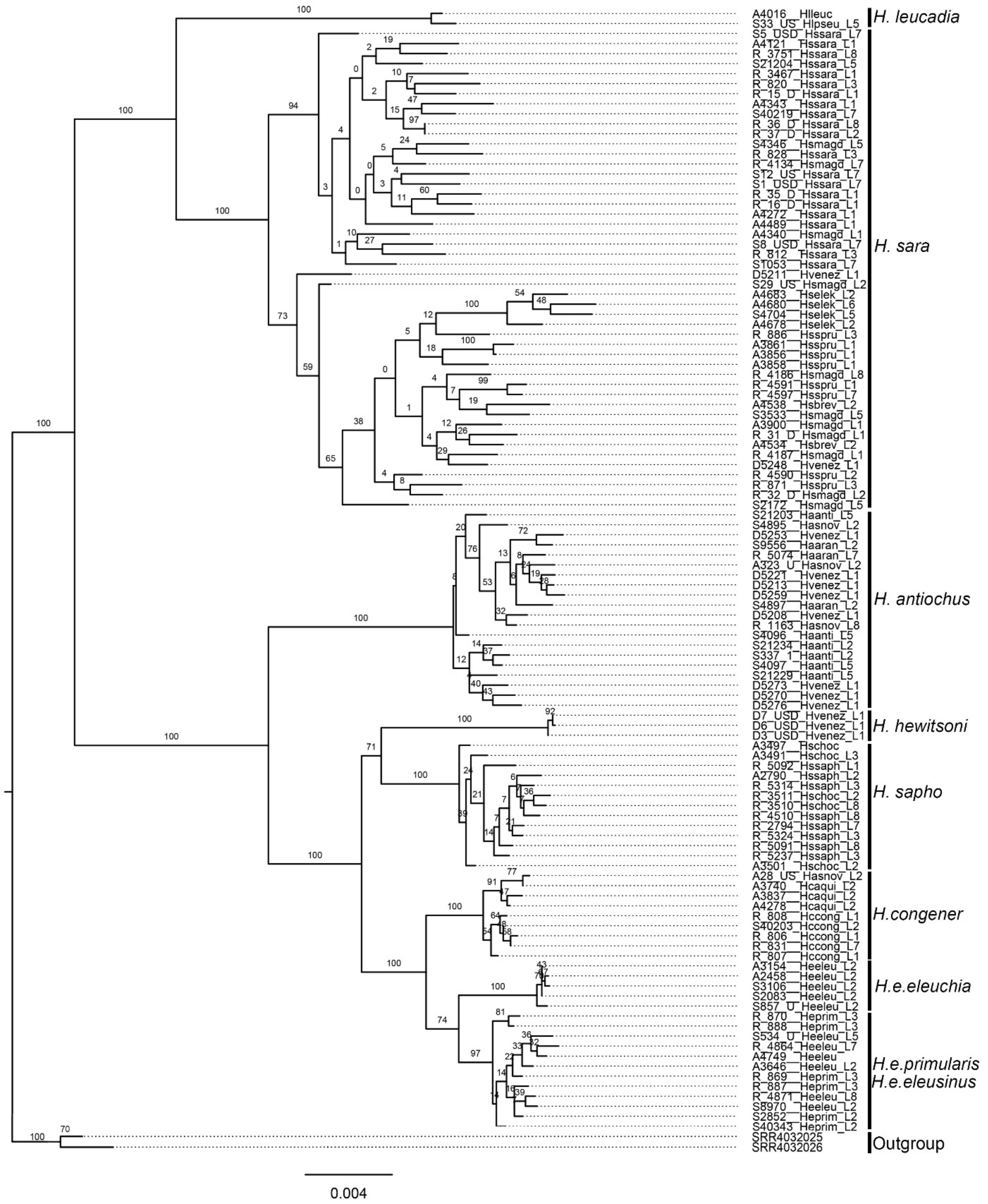


**Fig S20. Maximum likelihood phylogeny of chromosome 15.** Bootstrap support values are indicated at branches.

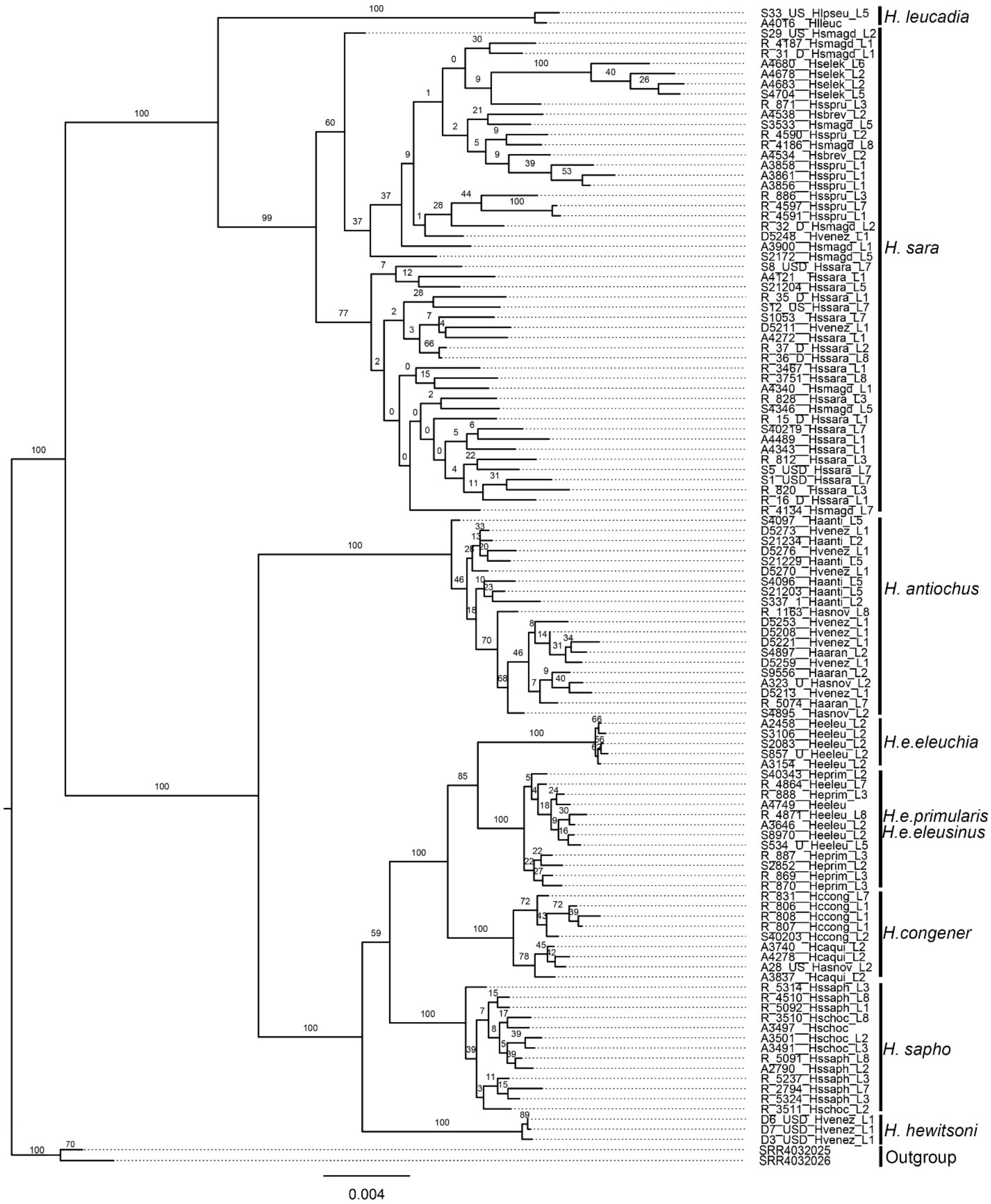


**Fig S21. Maximum likelihood phylogeny of chromosome 16.** Bootstrap support values are indicated at branches.

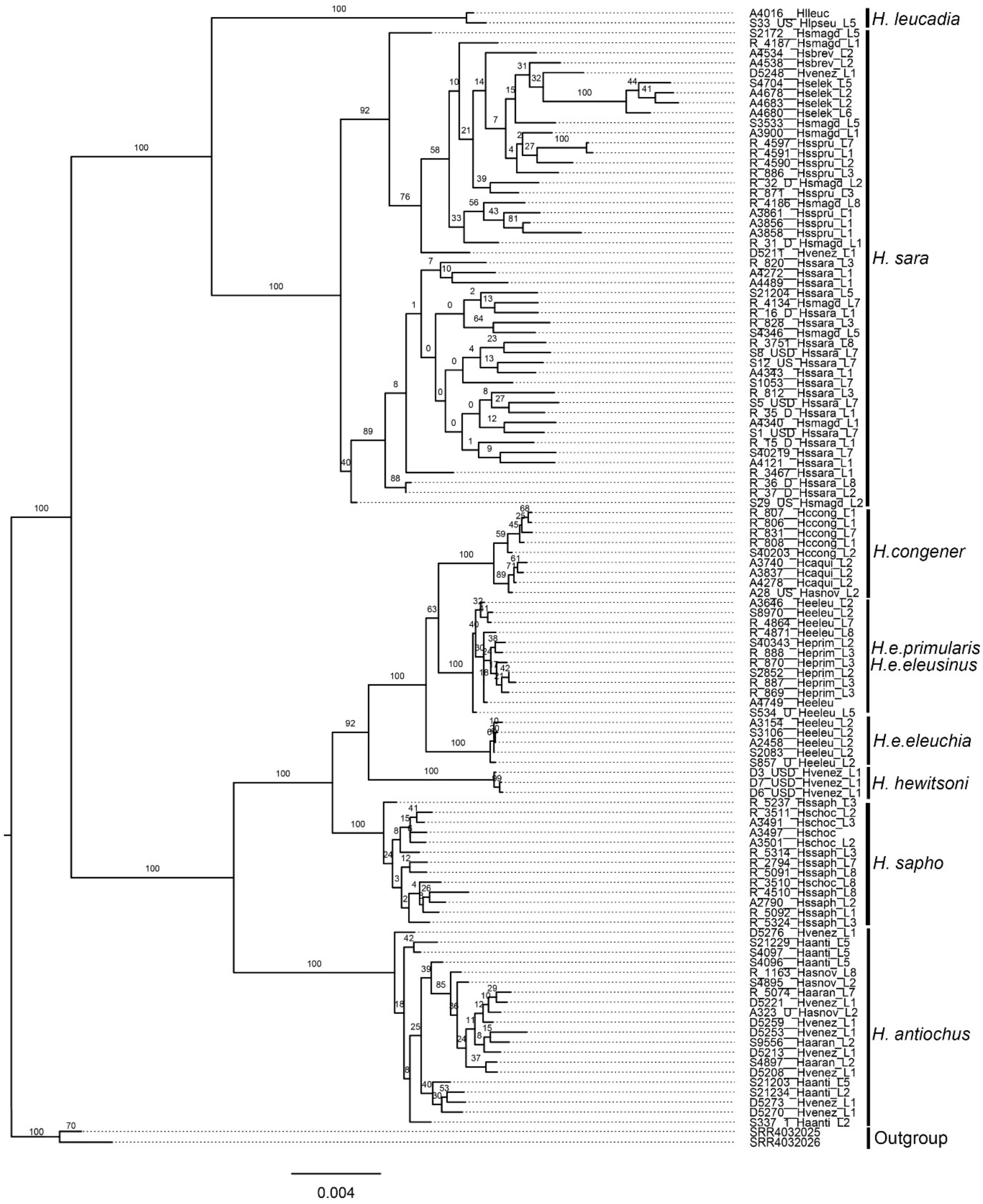


**Fig S22. Maximum likelihood phylogeny of chromosome 17.** Bootstrap support values are indicated at branches.

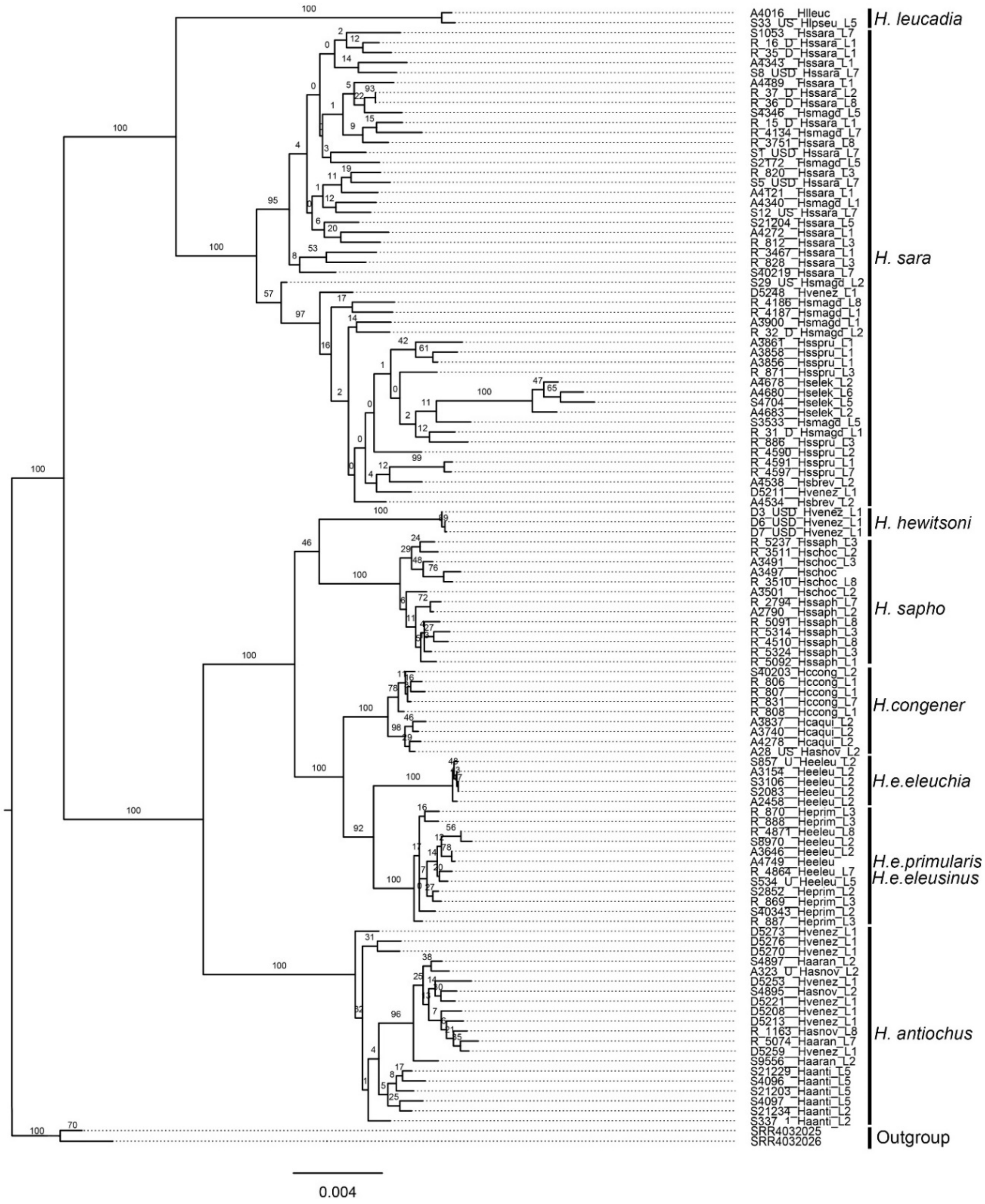

**Fig S23. Maximum likelihood phylogeny of chromosome 18 of the Sara-Sapho clade.** Bootstrap support values are indicated at branches, and the scale bar represents the percentage of substitutions per site.

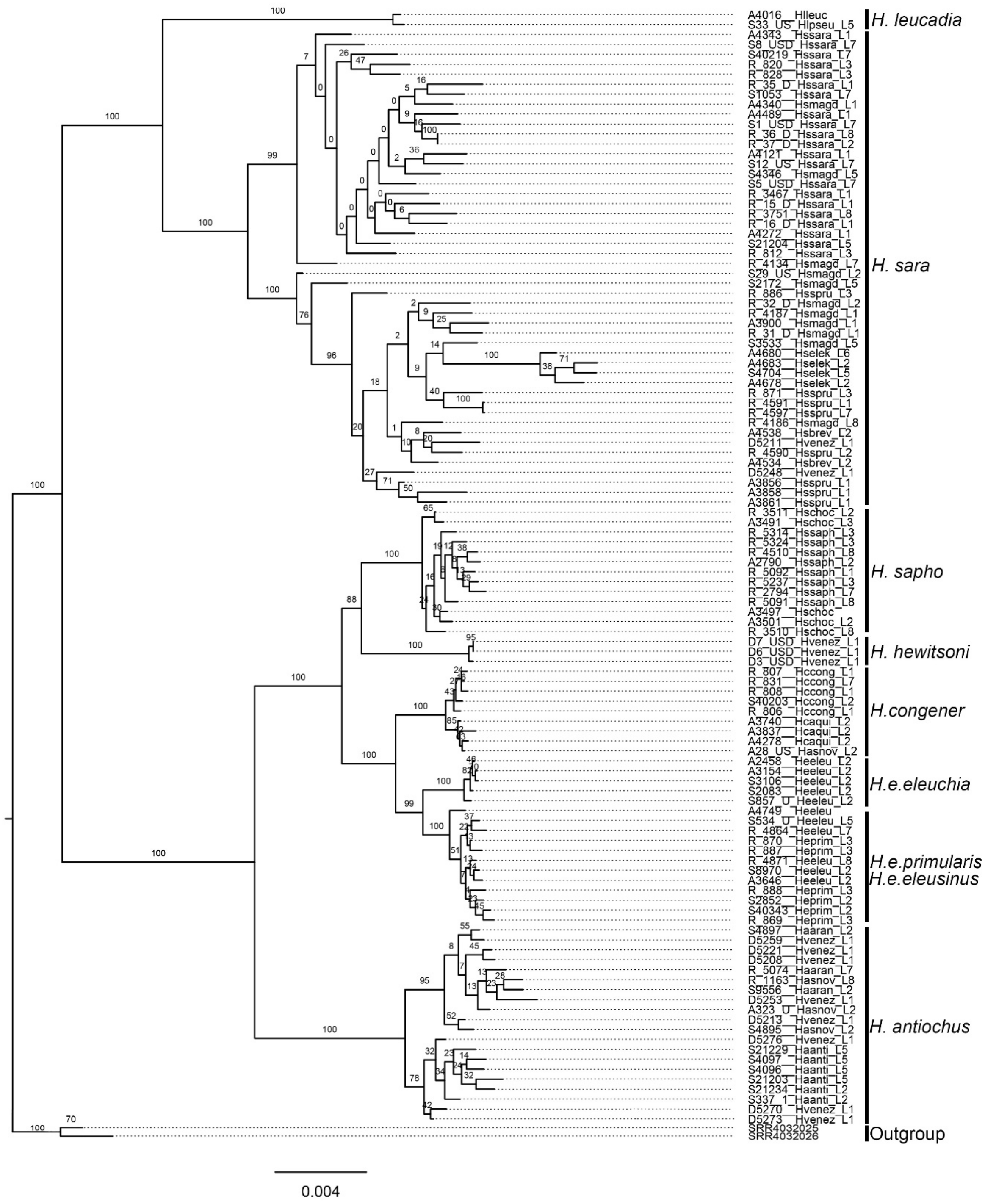


**Fig S24. Maximum likelihood phylogeny of chromosome 19.** Bootstrap support values are indicated at branches.

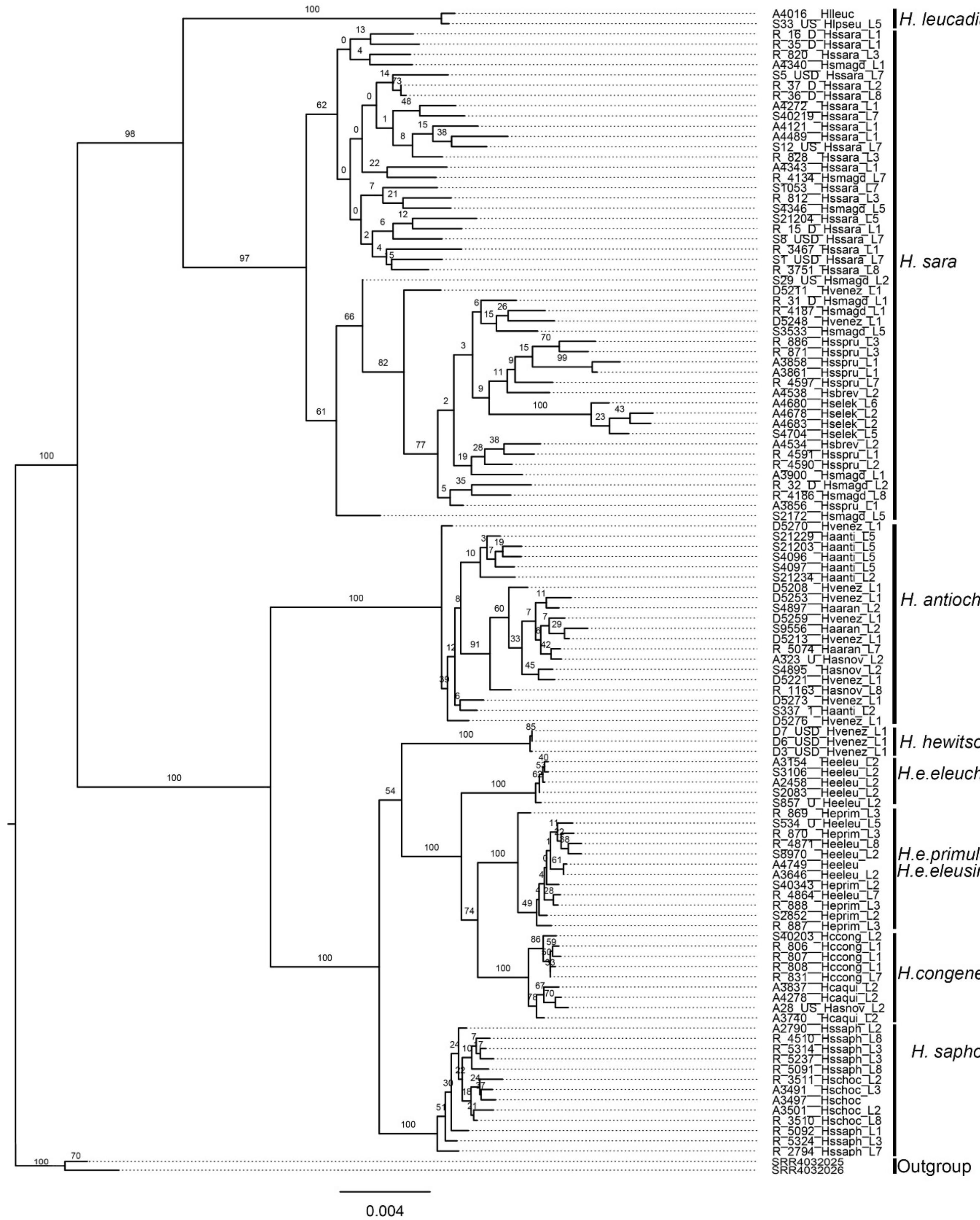


**Fig S25. Maximum likelihood phylogeny of chromosome 20.** Bootstrap support values are indicated at branches.

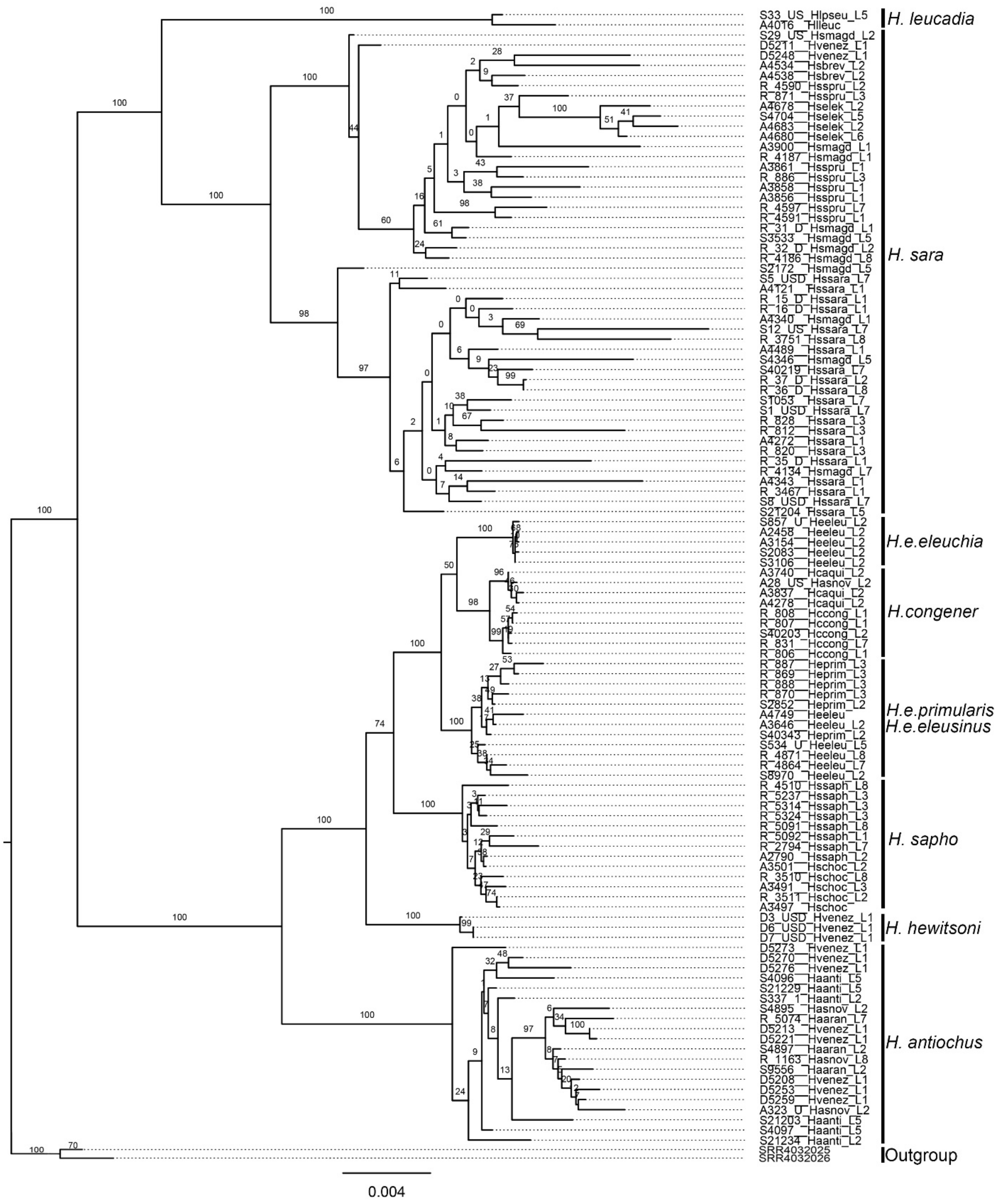


**Fig S26. Maximum likelihood phylogeny of chromosome 21.** Bootstrap support values are indicated at branches.

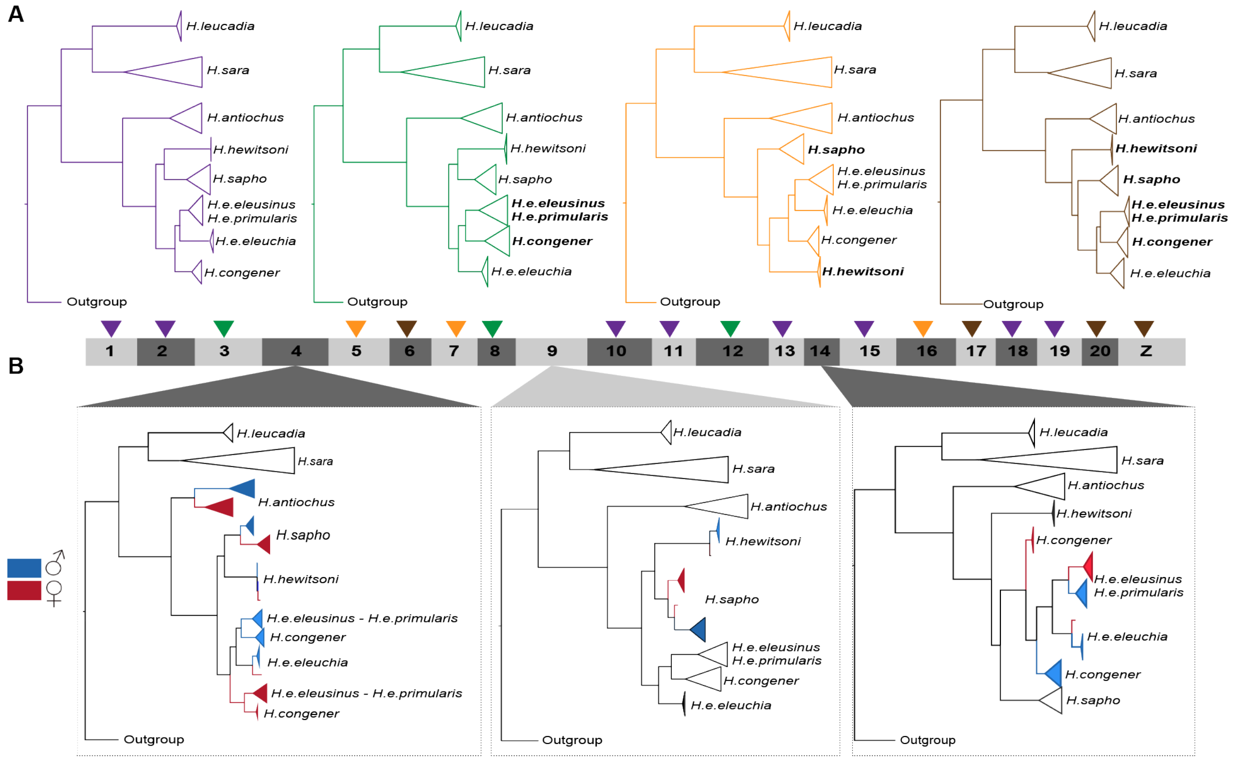


**Fig S27. Maximum Likelihood (ML) phylogenies inferred genome-wide and per chromosome.** (A) Topologies found across the genome. Purple: genome-wide topology. Green: *H. congener* within *H. eleuchia*. Orange: *H. hewitsoni* as sister to *H. congener* + *H. eleuchia*. Brown: *H. hewitsoni* as sister to *H. congener* + *H. eleuchia* + *H. sapho*. Chromosomes are shown in the bottom, with coloured triangles indicating the topology revealed by each of them. (B) Topologies showing sex-specific grouping within some species, which is indicative of Sex-A fusions in Chr4, Chr9 and Chr14. In these species, females are coloured in red and males in blue.

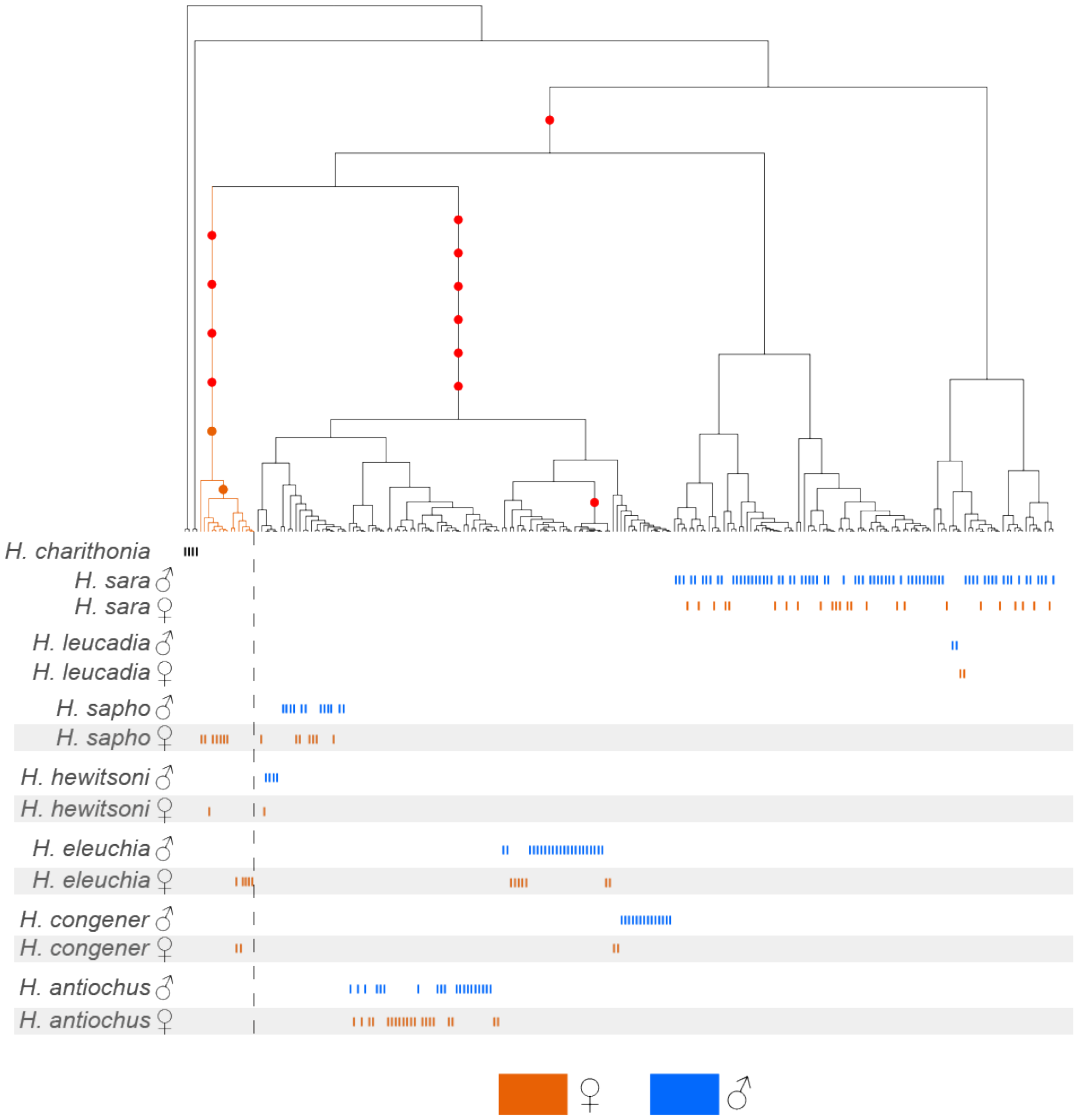


**Fig S28. Marginal tree for one SNPs in Chr4.** (A) This is one of the 112 SNP genealogies where female alleles from at least two species in the sapho subclade cluster as expected in a Sex-A fusion. Each vertical line represents an individual allele, and the alleles of all individuals are shown differentiating those of females (orange) from those of males (blue). Note that one allele of the females clusters with the alleles of males, while the other female allele formed a separate group (highlighted in orange).

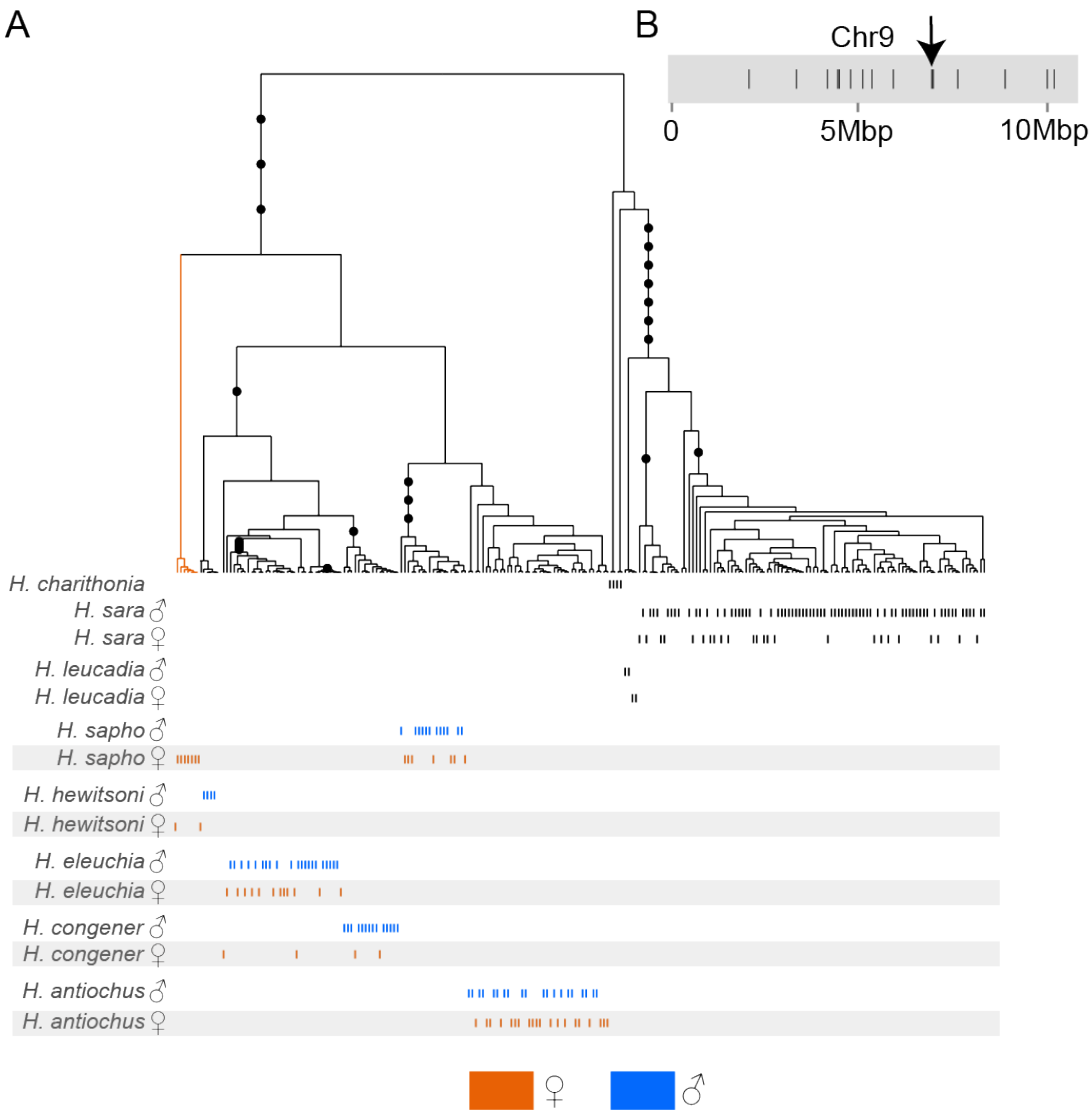


**Fig S29. Marginal tree for one SNPs in Chr9.** (A) This is one of the 15 SNP genealogies that showed a consistent pattern with the W-Sex fusion. The vertical lines represent the alleles of each SNP for each individual. Each vertical line represents an individual haplotype, and the alleles of all individuals are shown differentiating those of females (orange) from those of males (blue). Note that one haplotype of the females clusters with the alleles of males, while the other female allele formed a separate group (highlighted in orange). B) Position of each of the 15 SNPs in Chr9 that show genealogies consistent with a W-Sex fusion; they were not clustered in a specific region but rather distributed along the entire chromosome. The position of the SNP whose genealogy shown in A, is indicated by an arrow.

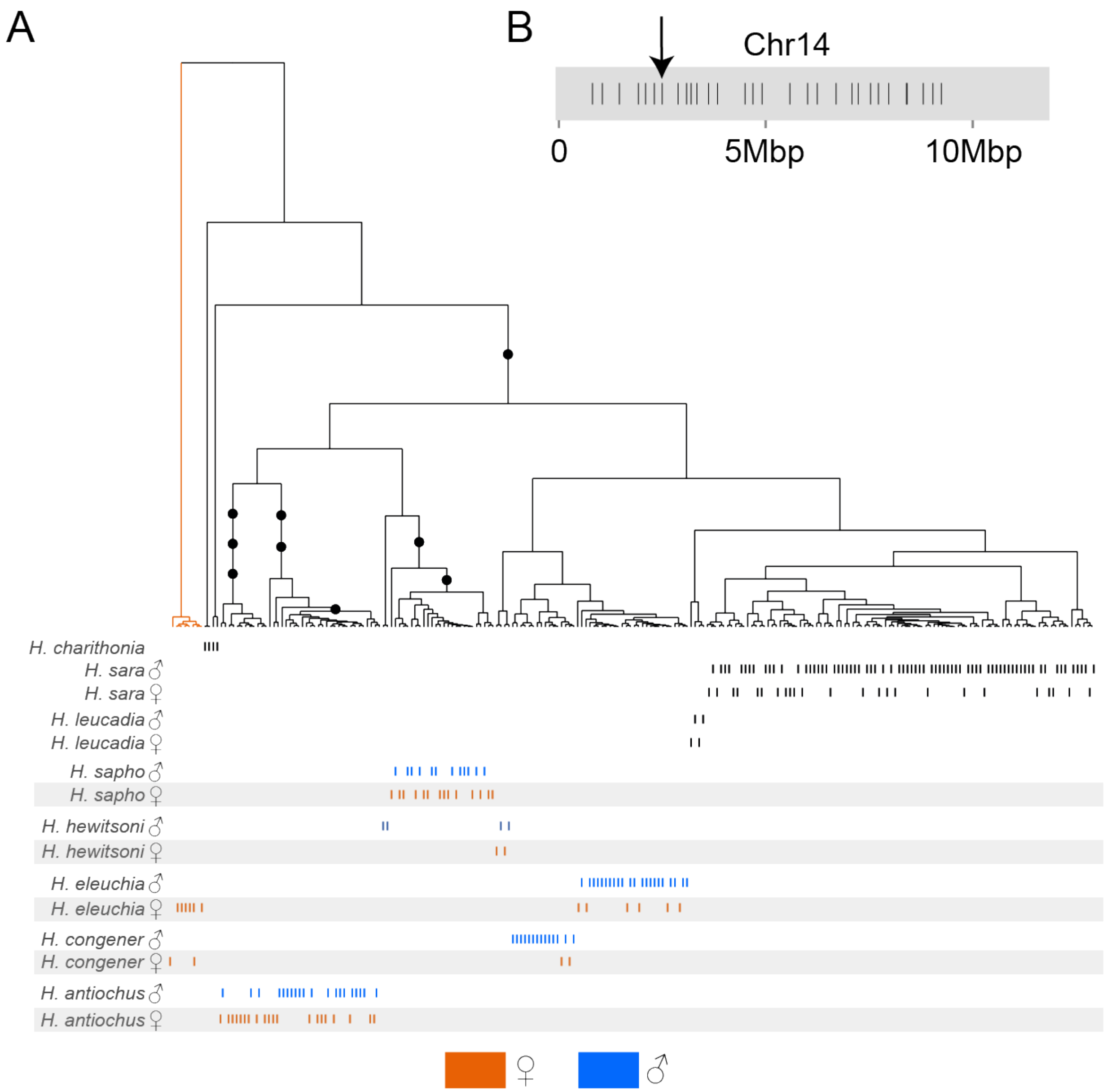


**Fig S30. Marginal tree for one SNPs in Chr14.** (A) This is one of the 25 SNP genealogies that showed a consistent pattern with the W-Sex fusion. Each vertical line represents an individual allele, and the alleles of all individuals are shown differentiating those of females (orange) from those of males (blue). Note that one allele of the females clusters with the alleles of males, while the other female allele formed a separate group (highlighted in orange). B) Position of each of the 25 SNPs in Chr14 that show genealogies consistent with a W-Sex fusion; they were not clustered in a specific region but rather distributed along the entire chromosome. The position of the SNP whose genealogy shown in A, is indicated by an arrow.

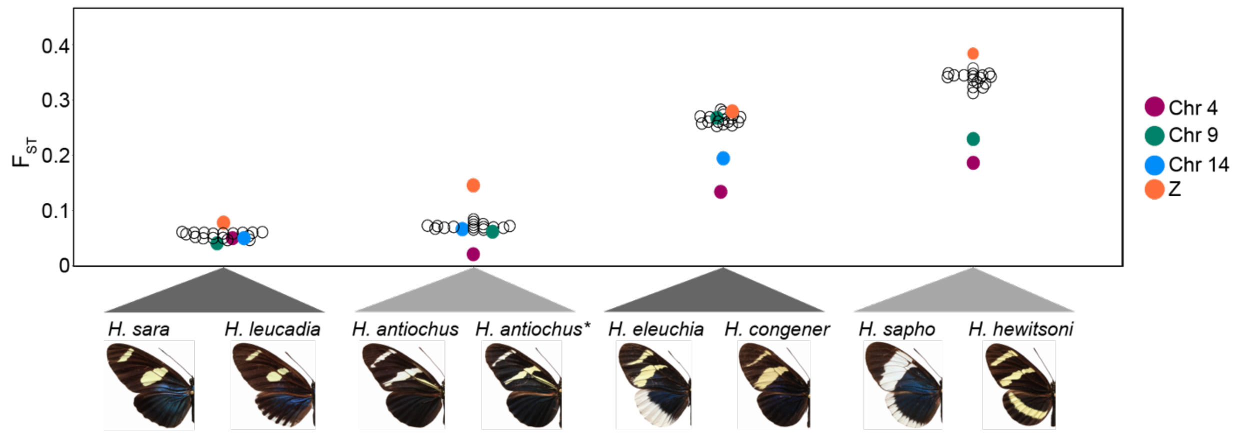


**Fig S31. Genome-wide divergence ($F_{ST}$) between pairs of species in the *sara/sapho* clade.** Each dot represents a chromosome, and chromosomes with evidence of Sex-A fusions are colour coded.

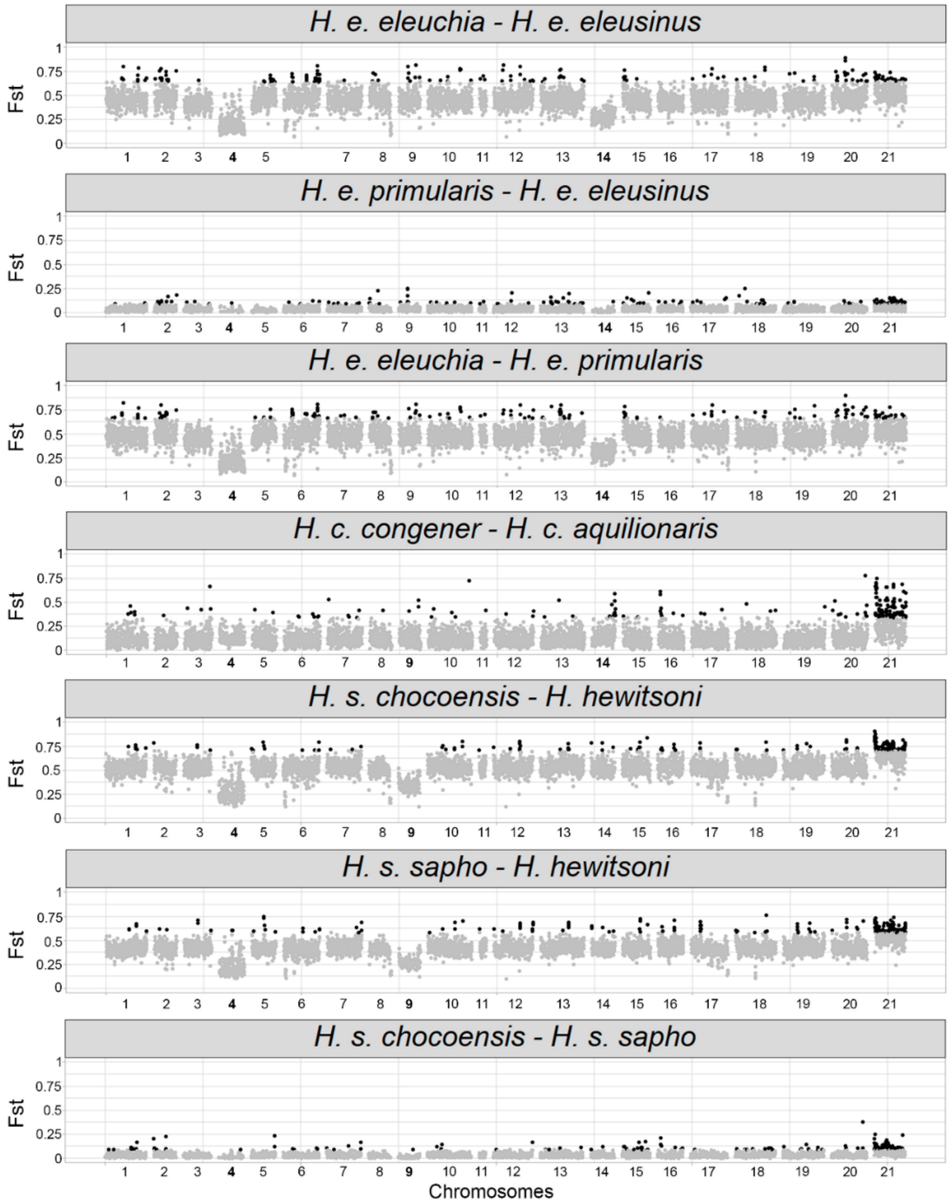


**Fig S32. Genome-wide divergence ($F_{ST}$) between pairs of subspecies of *H. eleuchia, H. congener* and *H. sapho*.** Each point represents a 50Kb window. The significance threshold is set at the top 5% of the $F_{ST}$ values distribution tail, and black windows are those that passed this threshold.

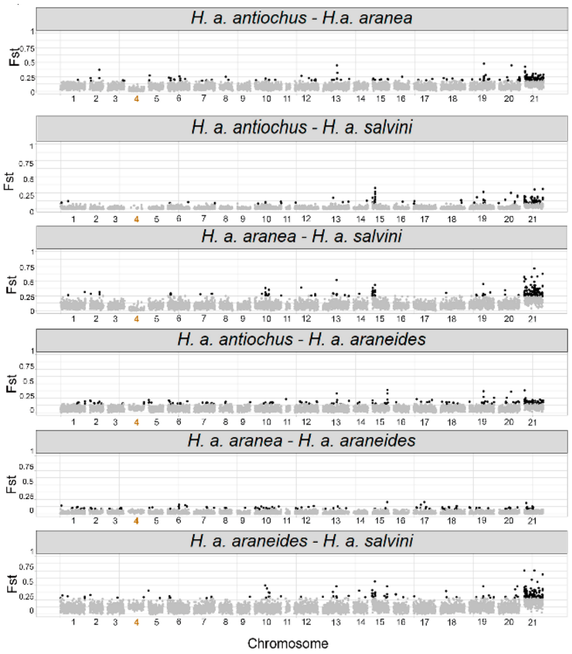


**Fig S33. Genome-wide divergence ($F_{ST}$) between pairs of subspecies of *H. antiochus*.** Each point represents a 50Kb window. The significance threshold is set at the top 5% of the $F_{ST}$ values distribution tail, and black windows are those that passed this threshold.

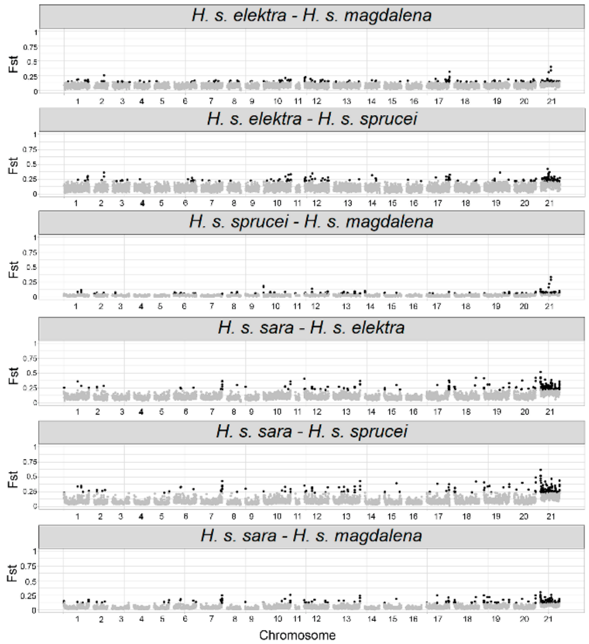


**Fig S34. Genome-wide divergence ($F_{ST}$) between pairs of subspecies of *H. sara*.** (a) $F_{ST}$. Each point represents a 50Kb window. The significance threshold is set at the top 5% of the $F_{ST}$ values distribution tail, and black windows are those that passed this threshold.

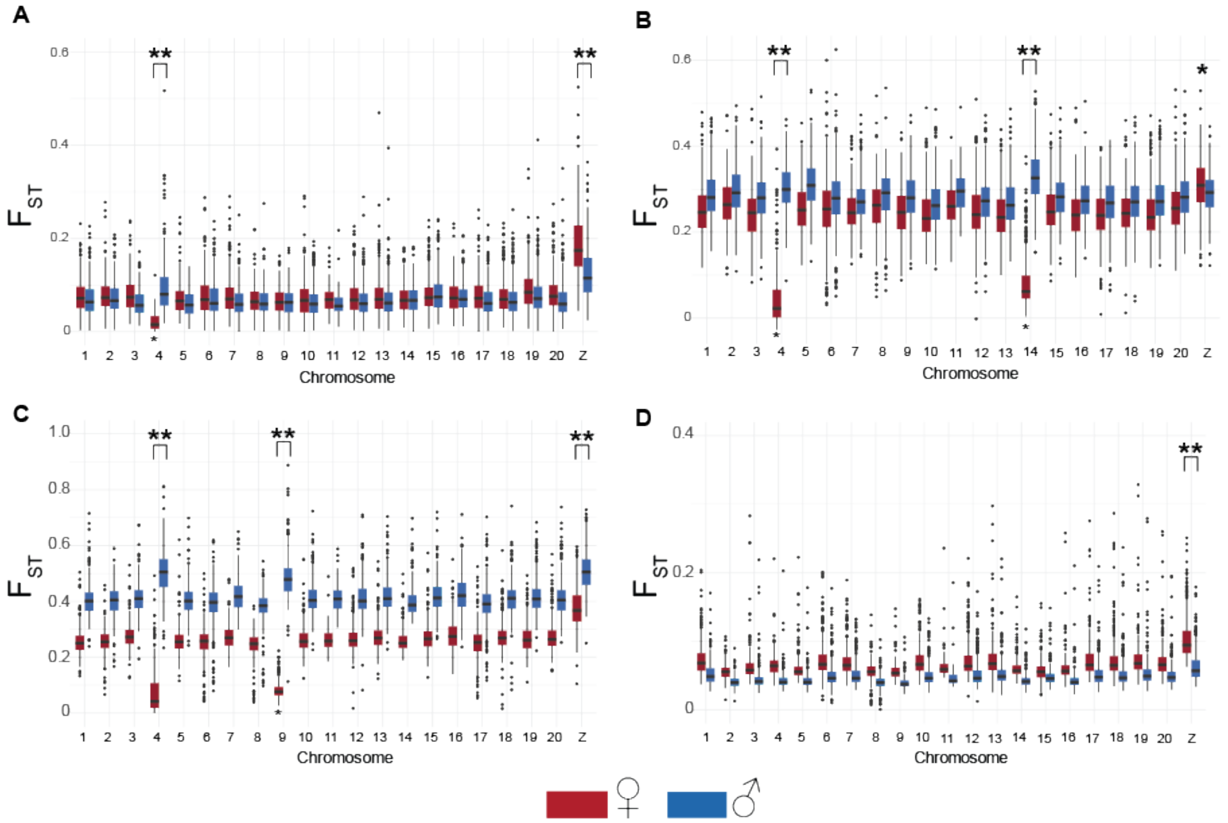


**Figure S35 Genome-wide divergence ($F_{ST}$) in the *sara/sapho* clade.** $F_{ST}$ between (A) subspecies of *H. antiochus*, (B) *H. congener* and *H. eleuchia*, (C) *H. sapho* and *H. hewitsoni*, and (D) *H. sara* and *H. leucadia*. $F_{ST}$ was calculated by sex and are colour coded. $^{**}p<0.05$.

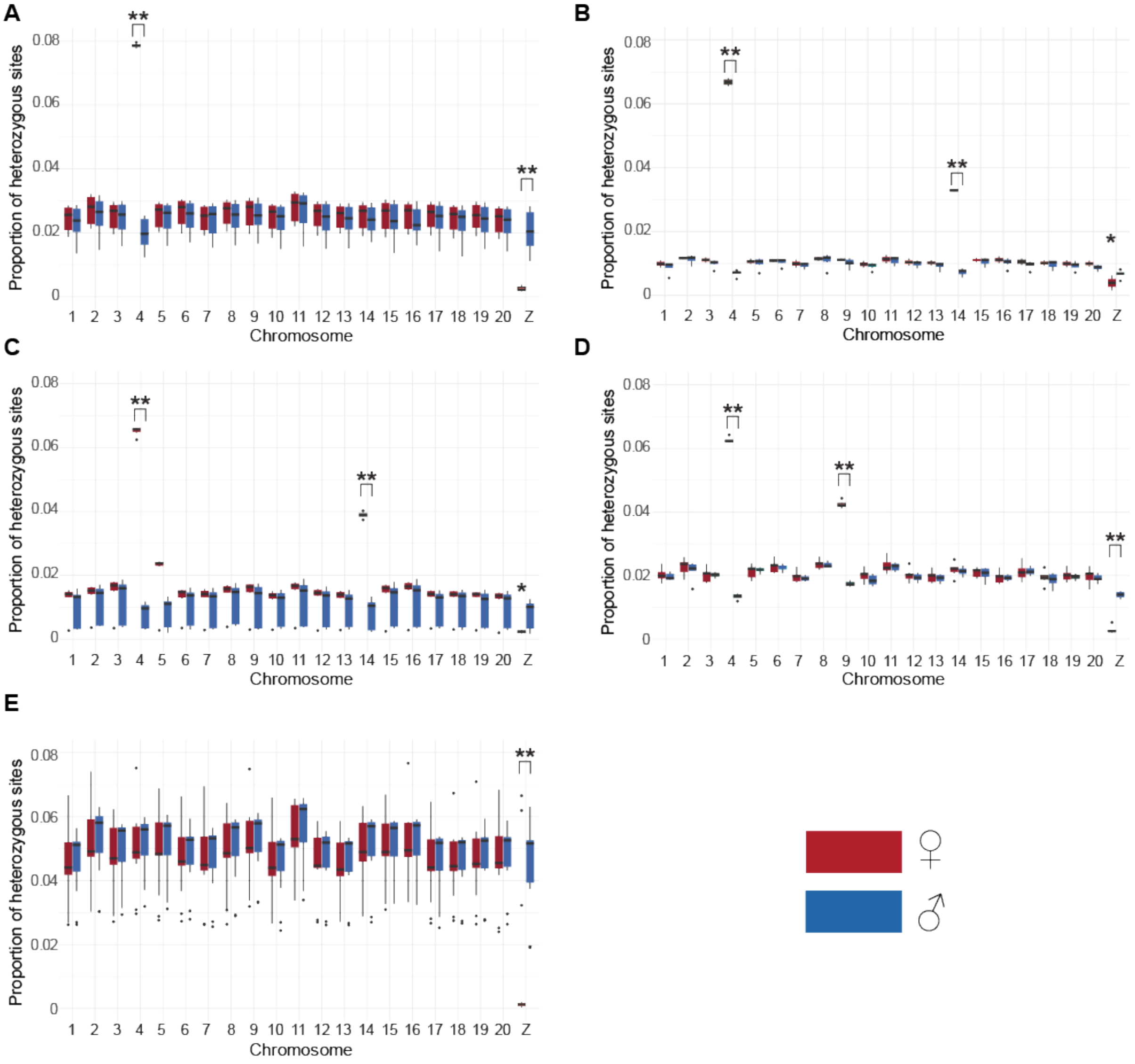


**Fig S36. Patterns of heterozygosity across the genome in the *sara/sapho* clade.** Proportion of heterozygous sites by sex in each species: (A) *H. antiochus*, (B) *H. congener*, (C) *H. eleuchia,* (D) *H. sapho*, and (E) *H. sara*. Sexes are colour coded. **p<0.05.

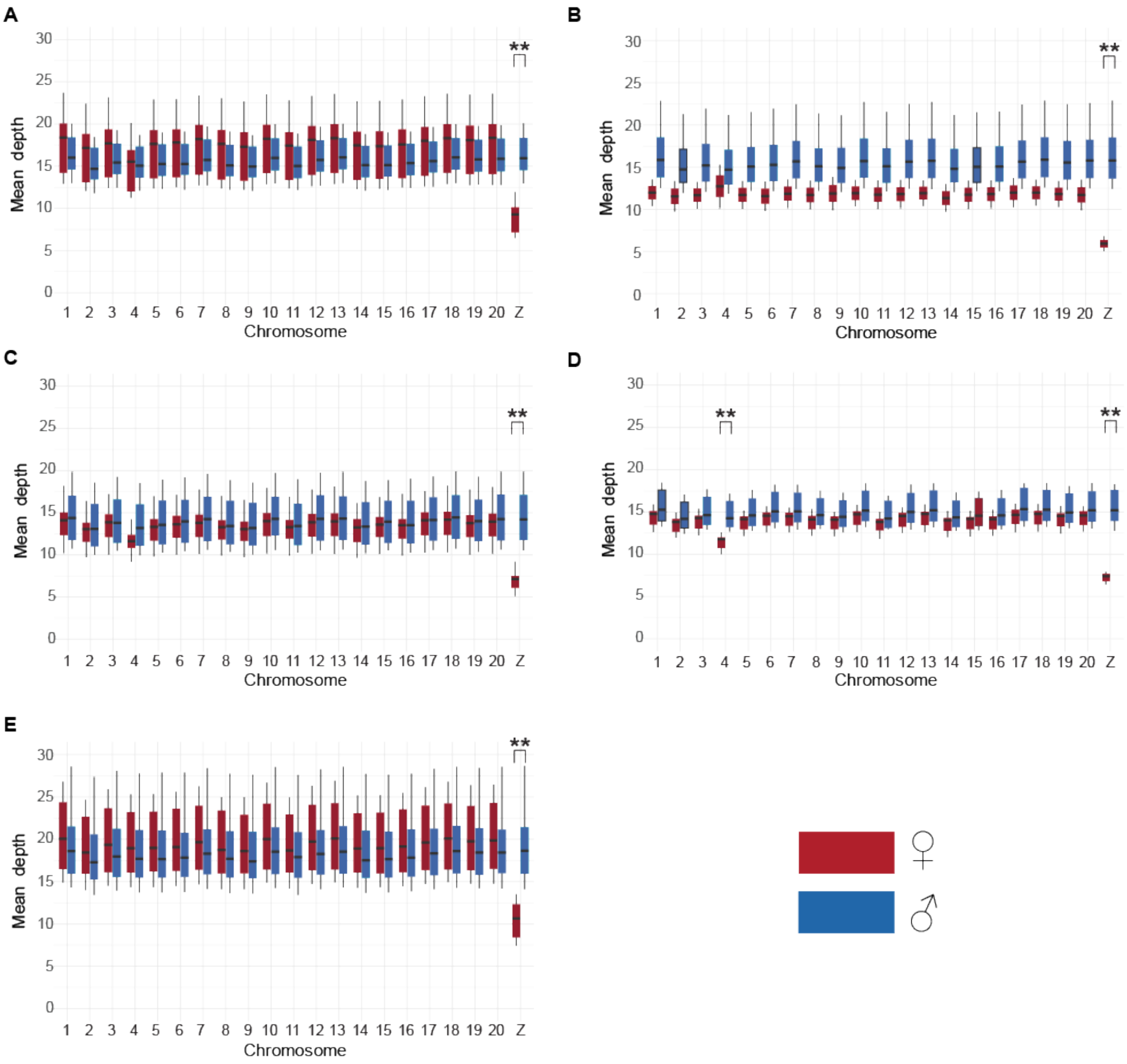


**Fig S37. Patterns of mean depth across the genome in the *sara/sapho* clade.** Mean depth by sex in each species: (A) *H. antiochus*, (B) *H. congener*, (C) *H. eleuchia,* (D) *H. sapho*, and (E) *H. sara*. Sexes are colour coded. **p<0.05.

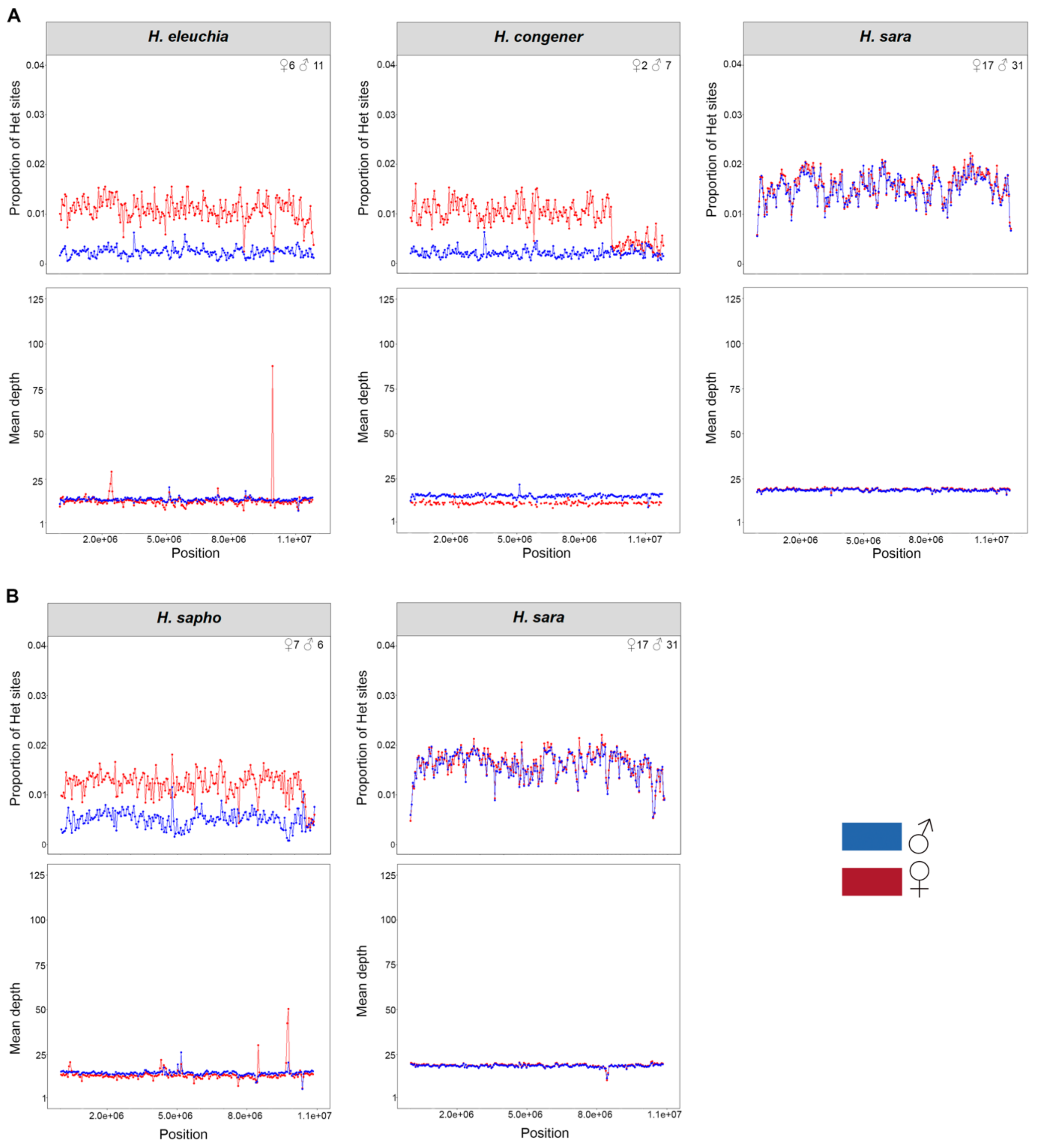


**Fig S38. Patterns of heterozygosity and depth across chromosome (A) 14 and (B) 9.** Proportion of heterozygosity sites and mean depth in 50Kb sliding windows in each species. Each line corresponds to one individual where males are shown in blue and females in red, and their *n* is shown in the top right corner. In all species, the sex-linked chromosomes show even patterns of increased heterozygosity, except for the end of Chr14 in *H. congener*.

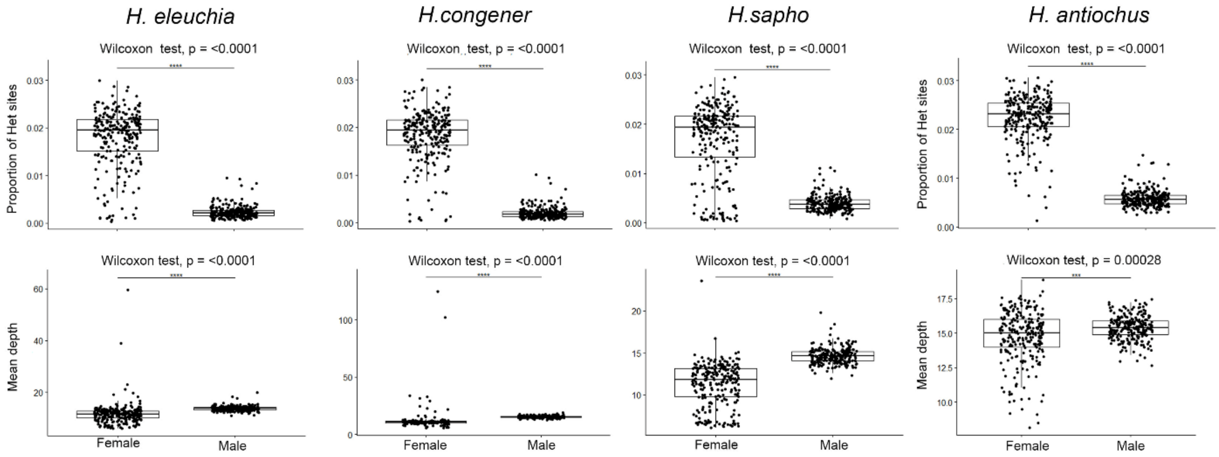


**Fig S39. Proportion of heterozygous sites and mean depth between sexes in sliding windows along chromosome 4.** Each panel corresponds to a species with the proportion of heterozygous sites shown at the top and mean depth at the bottom. Each dot represents the average of these values across all individuals per window. ns = non-significant.

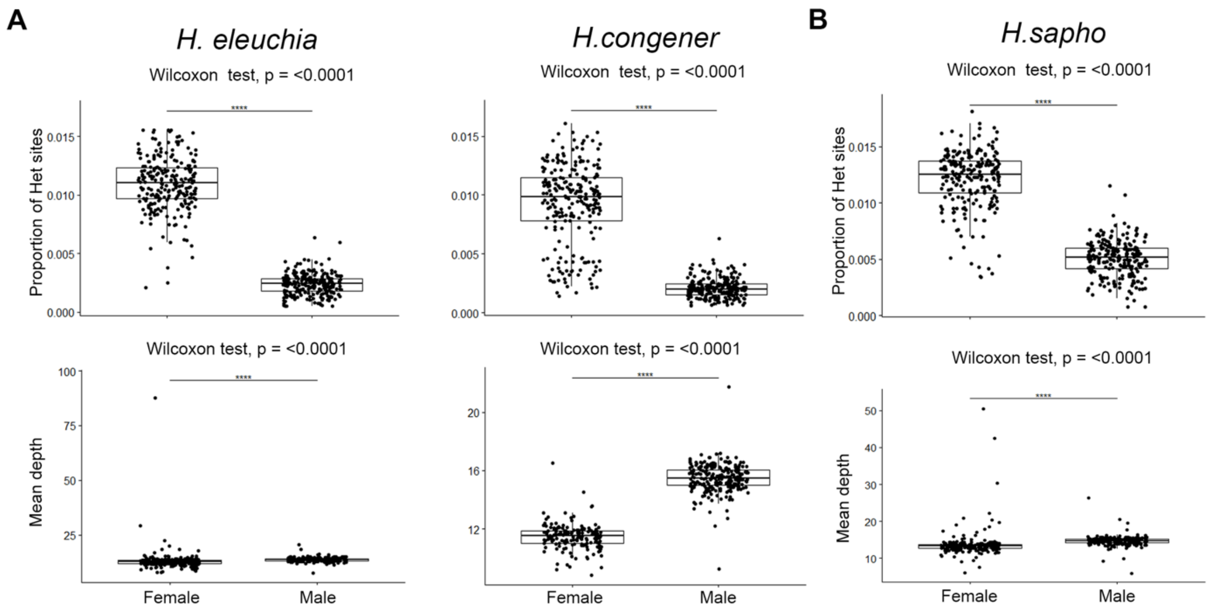


**Fig S40. Proportion of heterozygous sites and mean depth between sexes in sliding windows along chromosome (A) 14 and (B) 9.** Each panel corresponds to a species with the proportion of heterozygous sites shown at the top and mean depth at the bottom. Each dot represents the average of these values across all individuals per window. ns = non-significant.

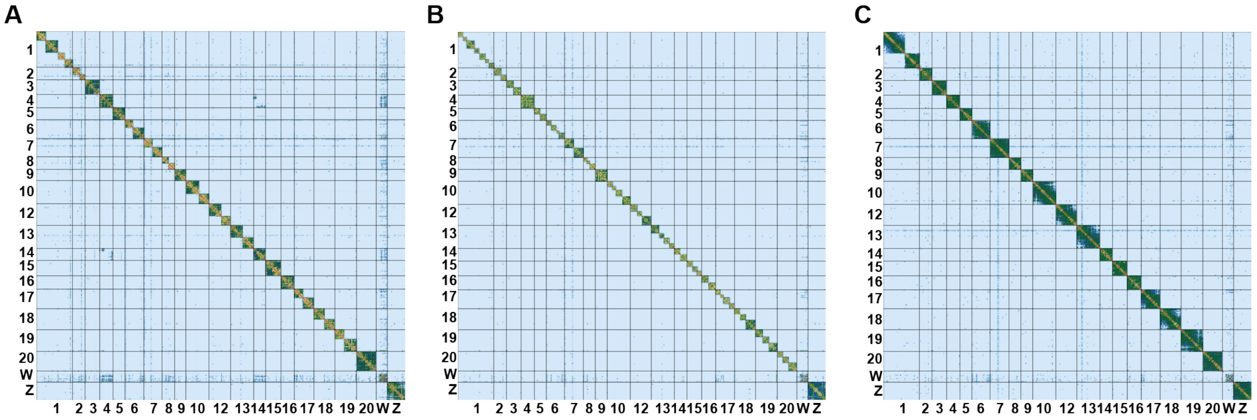

**Fig S41. The density map of Hi-C contacts confirms W-A fusions in the sapho subclade.** Hi-C contact heatmaps of the genome assembly of *H. charithonia* against: (A) the *H. sara* female (Hi-C used to assemble the genome reported here), (B) the *H. sapho* male, and (C) the female of *H. congener*.
